# Extrasynaptic NMDA receptors bidirectionally modulate intrinsic excitability of inhibitory neurons

**DOI:** 10.1101/2021.10.17.464733

**Authors:** Lulu Yao, Yi Rong, Xiaoyan Ma, Haifu Li, Di Deng, YongJun Chen, Sungchil Yang, Tao Peng, Tao Ye, Feixue Liang, Nenggui Xu, Qiang Zhou

## Abstract

The NMDA subtype glutamate receptors (NMDARs) play important roles in both physiological and pathological processes in the brain. Comparing to their critical roles in synaptic modifications and excitotoxicity in the excitatory neurons, much less is understood about the functional contributions of NMDARs to the inhibitory/GABAergic neurons. By using selective NMDAR inhibitors and potentiators, we here show that NMDARs bi-directionally modulate the intrinsic excitability (defined as spontaneous/evoked spiking activity and EPSP-spike coupling) in the inhibitory/GABAergic neurons. This modulation depends on GluN2C/2D-but not GluN2A/2B-containing NMDARs. We further show that NMDAR modulator EU1794-4 mostly enhances extrasynaptic NMDAR activity, and by using it we demonstrate a significant contribution of extrasynaptic NMDARs to the modulation of intrinsic excitability in the inhibitory neurons. Altogether, this bidirectional modulation of intrinsic excitability reveals a previously less appreciated importance of NMDARs in the second-to-second functioning of inhibitory/GABAergic neurons.

## Introduction

The NMDA subtype glutamate receptors (NMDARs) are important for both physiological and pathological processes during which their adequate activation is required for certain critical brain functions (such as learning and memory, and developmental refinement of neuronal connections), while their excessive activation during certain pathological conditions is believed to induce synapse loss and even neuronal loss ^1–4^. However, these conclusions are largely based on studies of NMDARs in the excitatory neurons, while the contribution of NMDARs in the inhibitory GABAergic neurons to brain physiology and pathology is far less understood ^5–7^.

Inhibitory neurons, although accounting for only about 20% neurons in the brain, play a key role in information processing, oscillations, tuning and proper balance between excitation and inhibition ^8, 9^. NMDARs in the inhibitory neurons have received much attention for their important contributions to pathology in many psychiatric disorders, including schizophrenia and depression ^2, 10–12^. However, detailed properties of the involved NMDARs, such as the contributions of synaptic and/or extrasynaptic NMDARs is poorly understood, comparing to the vast literature on the excitatory neurons ^12, 13^.

Previous studies have revealed the important contributions of NMDARs to synaptic plasticity and neuronal intrinsic properties, both are critical to the activity-dependent remodeling of neural circuit ^14–16^. Although the contribution of NMDARs to synaptic plasticity is widely studied and well-understood, how they contribute to the modification of the intrinsic neuronal excitability is less elucidated ^17, 18^. Intrinsic excitability, defined as the excitability in the absence of synaptic inputs, is determined by the expression of ion channels and receptors that contribute to the electrical properties of neurons ^19, 20^. To investigate this in the inhibitory neurons, we have used spontaneous spiking, injected current-induced spiking and EPSP-spike coupling (E-S coupling) as readouts ^21, 22^. Spontaneous spiking is essential for tuning the function of neuronal networks, while E-S coupling has significant contribution to the integration of synaptic excitation to generate action potentials in determining neuronal outputs^21^.

Subcellular localization and subunit composition are two important contributors to NMDAR functions, and both have been extensively explored in the context of synaptic plasticity and excitotoxicity ^1, 2, 10, 23^. It is well-established that NMDARs are located at both synaptic and extrasynaptic regions, with extrasynaptic NMDARs activated by spillover of synaptic glutamate or glutamate released from glia ^24, 25^, while synaptic NMDARs are activated by synaptically released glutamate but not extracellular glutamate. While the above conclusions are true for excitatory neurons, we recently showed that ambient glutamate can readily activate synaptic NMDARs in the inhibitory neurons due to the low presence of glutamate transporter GLT-1 ^26^. As for subunit composition, studies support a general conclusion that GluN2A and GluN2B subunits are highly expressed in the excitatory neurons while GluN2C and GluN2D subunits are more concentrated in the inhibitory GABAergic neurons ^2, 27, 28^. In addition, GluN2A-NMDARs are more likely to be concentrated at synapses, while GluN2B-NMDARs and GluN2C/2D-NMDARs are more enriched at extrasynaptic sites in the excitatory neurons ^10^. Limited evidences suggest that extrasynaptic NMDAR-mediated currents might contribute to the excitability of inhibitory neurons ^27, 29^. It needs to be noted that the contributions of NMDARs to physiology and pathology are likely more complex than a simple dichotomy between GluN2A- and GluN2B-NMDARs, or synaptic and extrasynaptic compartments. Thus, a more thorough understanding of the functional contribution of NMDARs on the inhibitory neurons to neuronal excitability will provide a deeper understanding of inhibitory neuron functions and the pathological processes involving them.

In this study, we used electrophysiological recording *in vitro* and *in vivo* to address how NMDARs may influence the excitability of inhibitory neurons, and the subunits and subcellular localization of participating NMDARs. We find bidirectional modulation of intrinsic excitability by NMDARs, which appears to be mediated at least partially by the GluN2C/2D-containing extrasynaptic NMDARs.

## Methods

### Animals

Mice were housed under standard conditions with free access to food and water. All experiments were carried out in accordance with the animal protection law and were approved by the Guangzhou University of Chinese Medicine, Peking University Shenzhen Graduate School Animal Care and Use Committee and the Animal Care and Use Committee at the Southern Medical University. Male and female mice of 6 -10 weeks, including wildtype (C57BL/6J), GAD67-GFP, PV-Cre, Ai14 and Ai32, and GluN2D-Flox transgenic mice were used. Mice were housed in a 12h light-dark cycle with free access to food and water.

### Electrophysiological recordings and analysis in vitro

Methods of brain slicing and patch-clamp recording were published previously ^26^. In brief, coronal frontal sections (400 μm) were cut on a VT1200S vibratome (Leica, Germany). Recording started at least 1hr after slicing on an upright microscope (Eclipse FN1, Nikon, Japan) at room temperature (23 ∼ 26 °C) with oxygenated aCSF (4 - 5 ml/min). Recordings were made from layer 2/3 PFC neurons, with GABAergic inhibitory neurons identified using GFP fluorescence in the GAD67-GFP transgenic mice. Signals were acquired at a sampling rate of 10 kHz and filtered at 2 kHz. To facilitate spontaneous spiking in the whole-cell configuration, most recorded neurons were held at − 45 ± 5 mV. For E-S coupling experiments, intensity electrical stimulation using a glass pipette was adjusted to trigger spikes with ∼ 50% probability for the first EPSPs. E-S coupling test using 5 pulses was given at 20 Hz unless mentioned otherwise.

NMDAR EPSCs were analyzed using Clampfit software (MDS Analytical Technologies). Average of NMDAR EPSCs in a 2 - 5 msec window centered on the maximal response was taken as peak amplitude. For measuring area, EPSCs were integrated from the start of responses to the time point with responses decayed to baseline (pre-stimulation) level. Stimulus artifacts were not subtracted in these measurements. For measuring the ambient NMDAR responses, slices were bathed in aCSF containing NBQX (10 μM), picrotoxin (50 μM) and low Mg^2+^ (0.5 mM), with neurons held at +40 mV. D-APV (100 μM) was added into the recording chamber via a thin tube positioned at the edge of the recording chamber.

To test whether GluN2C/2D-containing NMDARs contribute to EU1794-4 effect, changes in holding current at +40 mV was measured by applying EU1794-4 (300 μM) directly onto slices in the presence of Veh or NAB-14 (Figure 5g). To measure ambient responses at near physiological condition, we tested the contribution of GluN2C/2D-containing NMDARs to the extrasynaptic NMDARs-mediated ambient response, changes in holding current induced by NAB-14/Pip18/DMSO were recorded at -60 mV. In order to amplify signals, slices were pre- bathed with NMDA (3 μM) (Figure 5i). All the response normalized to that application of D-APV. To examine the potential activity-dependent property of EU1794-4, baseline NMDAR mediated synaptic response was acquired using 1 Hz stimulation, and 20 Hz stimulation (for 50 sec) was used after sufficient time was allowed for EU1794-4 to take effect.

### Western blot

The GluN2D-floxp transgenic mice were injected with vGAT-cre virus (BrainVTA, Wuhan) into the PFC (AP: 1.94 mm anterior to Bregma; M/L: ±0.4 mm lateral to midline; D/V: 2.65 ventral from the cortical surface). 21d expression after virus injection, mice were anesthetized and perfused with phosphate-buffered saline (PBS) through the heart. Freshly dissected mouse brains were treated with RIPA lysis and homogenized by S10-High Speed Homogenizer (Xinzhi Biotechnology Co., Ltd., China), and then centrifuged at 17,925 × g for 10 min at 4 °C. Protein concentration was measured with a BCA Protein Assay Kit (Pierce). Clarified cell extracts were mixed with 6× sodium dodecyl sulfate (SDS) sample buffer. Protein samples were run on 8% SDS-polyacrylamide gel electrophoresis using a Bio-Rad gel system and transferred onto nitrocellulose membranes. Loading controls (glyceraldehyde 3-phosphate dehydrogenase (GAPDH)) were run on the same gel. Membranes were then probed with antibodies with the appropriate dilutions, including anti-GLT-1 (1:2000; Abcam, catalog no. ab178401) and anti-GADPH (1: 10,000; Sigma, catalog no. G8795). ImageJ was used for densitometric analysis. Experiments were performed in a double-blind manner.

### *In vivo* single cell recording

Male and female wild type (C57BL/6J) and transgenic (PV-Cre, Ai14 and Ai32) mice aged 2-3 months were used. The method was described in the previous studies ^30–32^. Mice were anesthetized with isoflurane (1.5%, v/v) and a head post for fixation was mounted on top of the skull with dental cement. Skull over the A1 was cleaned and protected from being covered by dental cement as we described previously ^30–32^. Afterward the mouse was injected subcutaneously with 0.1 mg kg-1 buprenorphine and returned to its home cage. During the recovery period, the mouse was trained to get accustomed to the head fixation on the recording setup. To fix the head, a screw was tightly clamped by a custom-made post holder. Recordings were performed in a sound attenuation booth. For recording, the mouse was briefly anesthetized with isoflurane and a craniotomy was performed over the A1. Meanwhile, the dura was removed to make the drug seep into A1. The mouse was then left to recover from isoflurane for at least 2 h. Recording experiments were started after the animal exhibited normal running behavior. Each recording session lasted for about 4 hr. The mouse was given drops of 5% sucrose (wt/vol) through a pipette every hour. Between sessions, the animal was returned to its home cage for a break of at least 2 hr.

Software for sound stimulation and data acquisition was custom made in LabVIEW (National Instruments). Stimuli include pure tones (2–64 kHz spaced at 0.1 octave, 50-ms duration, 3-ms ramp, 0 - 70dB sound pressure level (SPL) spaced at 10 dB, in pseudorandom sequence, 3 repetitions, 0.5-s interstimulus interval), CF and noise (Characterize Frequency and broadband white noise, 50-ms duration, 3-ms ramp, 60 dB SPL, 1s interstimulus interval). The loudspeaker (Brüel and Kjaer 4138, 4135, Naerum, Denmark) and an amplifier (Brüel and Kjaer 2610, Naerum, Denmark), was placed 10 cm away from the animal’s head facing the unsealed ear.

Neurons were recorded in heterozygous offspring of a cross between a cre-dependent ChR2-EYFP (Ai32) line and a PV-tomato-Cre line (heterozygous offspring of a cross between PV-Cre line and Ai14 line). Loose patch recordings using pipettes of smaller tip openings (pipette impedance, ∼10 MΩ) were performed. An optic fiber and drug infusion tube were positioned close to the cortical surface of the recording site. PV-neurons were identified with 10 pulses (50 ms duration, 50 ms interstimulus interval) of blue LED stimulation (470 nm, Thorlabs). PV-neurons responded to 50 ms light pulses with a reliable, short-latency burst of spikes and showed sustained spiking responses.

To verify the PV+ expression, animals were deeply anesthetized with urethane (25%) and transcardially perfused with 4% paraformaldehyde (PFA) in phosphate-buffered saline (PBS). The brain tissue was sliced into coronal sections (150-μm thickness) by using a vibratome (Leica Microsystems). Slices were imaged using a confocal microscope (Olympus FluoView FV1000).

We performed data analysis with custom-developed software (MATLAB, MathWorks). Tone- and Noise-evoked spikes were counted within a 100 ms window after the onset of stimuli. Evoked firing rate (FR) was calculated after subtracting the average baseline FR. Evoked responses were defined as firing rates higher than the average baseline firing rate by 3 standard deviations. Neurons that did not exhibit evoked spiking responses were excluded from the analysis.

### Multi-channel single unit recording

Experimental methods were described previously ^33^. Briefly, mice were anesthetized with isoflurane (∼ 1% in a gas mixture) on a stereotaxic apparatus (RWD Life Science Co., China). A craniotomy was made over the lateral ventricle (AP: − 0.5; ML: − 1.0; DV: 2.0). A hole was drilled through the skull. Infusion kit was slowly lowered with a gripper, until the pedestal touched the skull. Electrode array was unilaterally implanted aiming at medial prefrontal cortex (mPFC) (1.94 mm anterior to Bregma; 0.4 mm lateral to midline; 2.65 ventral from the cortical surface), and sealed to the skull with dental cement. After surgery, mice were allowed to recovery for 7 - 14 days. During recording of spontaneous spiking, mice moved freely inside a recording box (390*180*180 mm). Baseline of 30 min was collected before injection of M-8324 (100 μM, 5 μL, i.c.v.), and recording was continued for 120 min afterwards. Individual recorded neurons were sorted using Plexon Offline Sorter and analyzed using Neuro Explorer (Nex Technologies, USA). Classification of excitatory and inhibitory neurons, calculation of E/I ratio were performed as described previously ^33^.

### Reagents

D-APV, NBQX and QX-314 chloride were obtained from Tocris; Picrotoxin, EU1794-4, NMDA were from Sigma-Aldrich; NAB-14, EU1794-4-Fluo and M-8324 were synthesized at Peking University Shenzhen Graduate School. Pip18 was provided by Genentech (South San Francisco, CA, USA).

## Results

### NMDARs bidirectionally modulate the intrinsic excitability of inhibitory neurons in vitro

Previous studies have suggested that NMDARs may contribute to the excitability of inhibitory neurons ^34–37^. To test whether this is the case, we examined the effect of bath applied NMDAR modulators on genetically identified GABAergic neurons in PFC slices from GAD-67 GFP mice. The spontaneous spiking rate was used for measuring excitability ^38^, and normal aCSF Mg^2+^ (1 mM) was used to mimic physiological conditions. We first tested the bath perfusion of a competitive NMDAR antagonist D-APV (100 μM), and observed reversibly reduced spontaneous spiking in most of the recorded GABAergic neurons (Fig. 1a, b). We then examined the impact of enhancing NMDAR activity using a NMDAR-PAM M-8324 which we have shown previously to selectively enhance the activity of NMDARs on inhibitory neurons (Deng, 2020). Bath perfusion of M-8324 (30 μM) led to significantly enhanced spontaneous spiking in the GABAergic neurons (Fig. 1c, d), which occurred to a similar degree in both FS and non-FS (RS, regular spiking) GABAergic neurons (Fig. 1e), suggesting that this modulation by NMDARs is bi-directional. This impact of M-8324 on excitability was not affected by GABAergic synaptic transmission (Sup Fig. 1).

**Figure 1.**
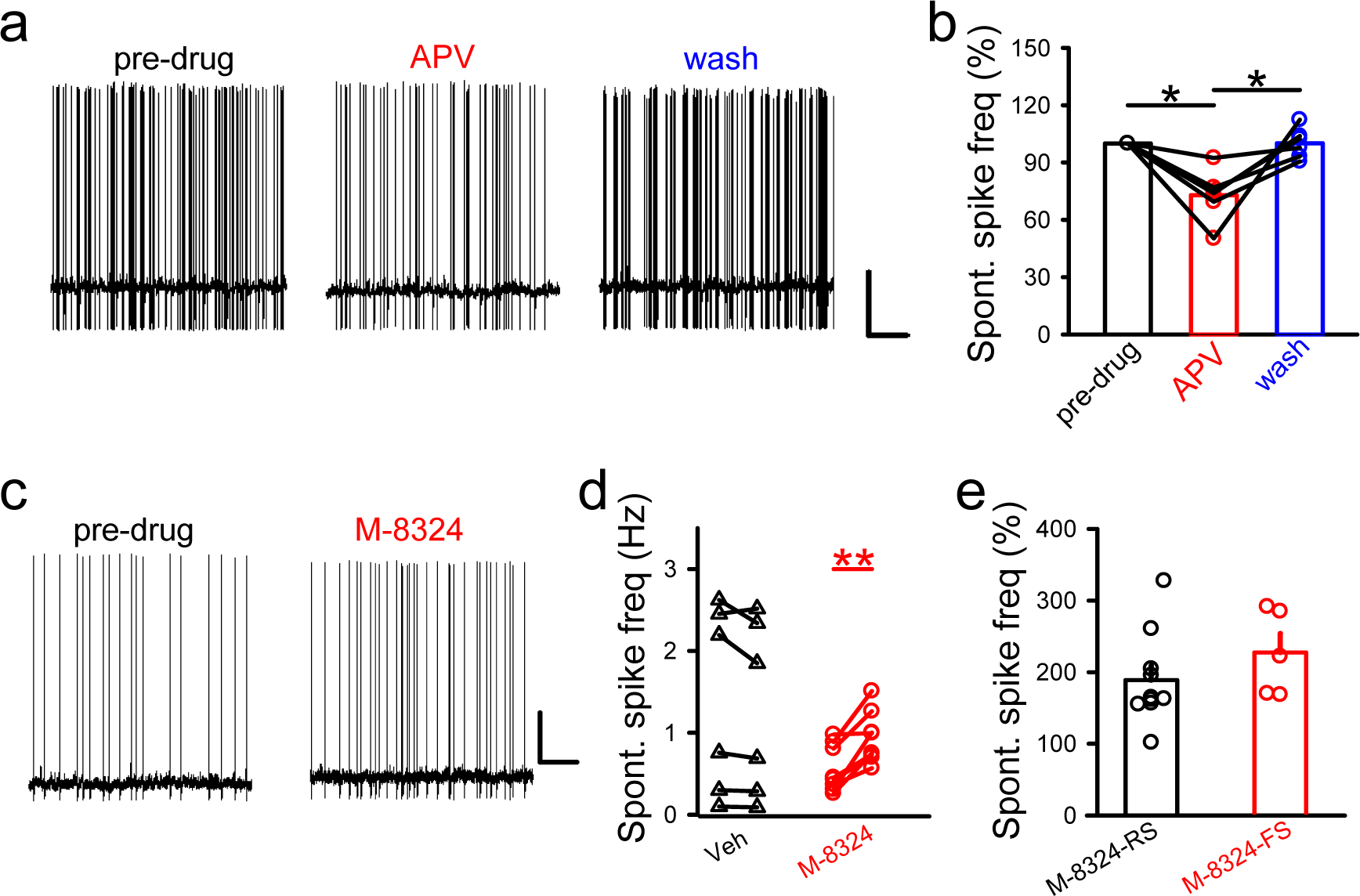
Bi-directional modulation of spontaneous spiking in PFC GABAergic/inhibitory neurons by NMDAR modulators. **a,** Sample traces of spontaneous spiking in identified GABAergic/inhibitory neurons before, during and after bath application of D-APV (100 μM) in PFC slices. Scale bars, 0.5 s/20 mV. **b**, Spontaneous spiking frequency was reversibly reduced by bath application of D-APV. APV, 72.92% ± 5.55; wash, 100.03% ± 3.23. N(cells)=6. *, *P*<0.05, paired t-test. **c,** Sample traces showing the impact of M-8324 (30 μM) on spontaneous spiking in GABAergic neurons. Scale bars, 5 s/20 mV. **d,** Quantification of M-8324 impact on spontaneous spiking in GABAergic neurons. Spike rate was normalized to the pre-drug level. Veh, 90.26% ± 2.82; M-8324, 190.75% ± 27.13. N(cells)=9 (Veh), N(cells)=8 (M-8324), * *P*<0.05, paired t-test. **e**, Changes in spontaneous spiking frequency in regular spiking (RS) and fast spiking (FS) GABAergic neurons. M-8324 (RS), 188.94% ± 20.11, N(cells)=10; M-8324 (FS), 227.35% ± 26.62, N(cells)=5. Data was shown as mean±SEM.

The above results suggest that NMDARs may modulate the intrinsic excitability of inhibitory neurons. Another method to measure the intrinsic excitability is to inject depolarizing current through the recording patch pipette. Significantly increased spike frequency was seen in the presence of M-8324 in the GABAergic neurons compared to that treated with vehicle (Fig. 2a), and to a similar degree in FS- and RS-neurons (Fig. 2b). In addition, spike frequency induced by current injection was significantly and reversibly reduced in the presence of D-APV (Fig. 2c). Thus, NMDAR modulation of current-induced spiking in the GABAergic neurons is also bi-directional. The spiking threshold, as measured by the amount of current needed to evoke a single spike, was significantly reduced in the presence of M-8324 (Fig. 2d), supporting higher excitability. To better mimic the natural trigger of spikes, we used depolarizing ramp current and observed that M-8324 significantly enhanced the probability of spiking (Fig. 2e). To determine whether changes in the passive properties of neurons might contribute, we measured input resistance (Rin), spike height, afterhyperpolarization (AHP), or half width of spike (Sup Fig. 2), and found none of them were altered by M-8324. Taken together, altered intrinsic excitability is likely a major contributor to the NMDAR modulators-induced changes in spiking.

**Figure 2.**
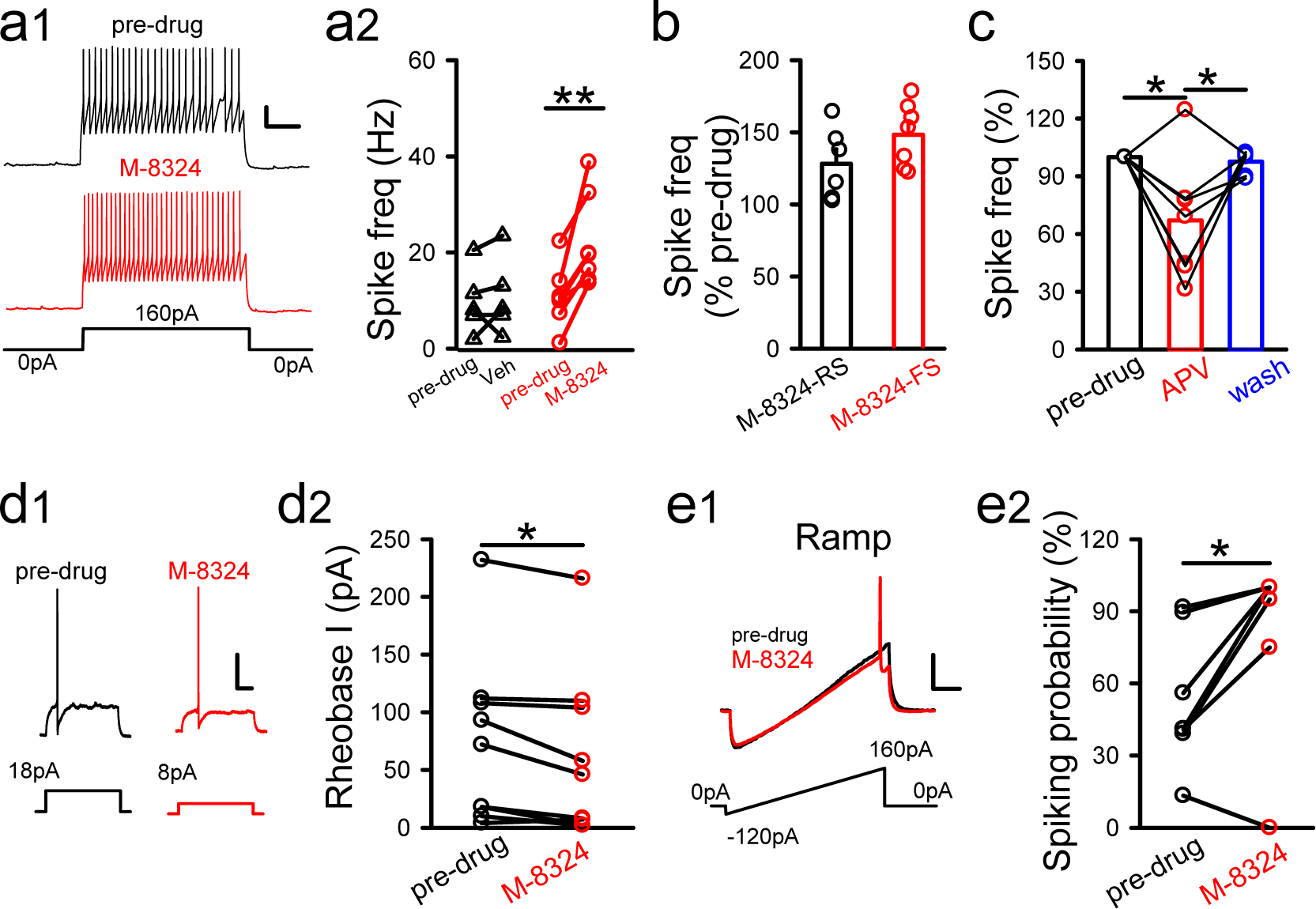
Impact of M-8324 on spikes triggered by current injections. **a1,** Sample traces showing M-8324 impact on spiking triggered by current injections in GABAergic neurons. Scale bars, 200 ms/20 mV. **a2,** Quantification of M-8324 impact. Pre-drug, 9.81 Hz ± 3.07; Veh, 10.87 Hz ± 3.58. N(cells)=7, * *P*<0.05, paired t-test **b,** M-8324 impact on spike frequency in RS- and FS-GABAergic neurons. M-8324 (RS), 128.19% ± 10.13, N(cells)=6; M-8324 (FS), 148.39% ± 8.34, N(cells)=7. **c,** Impact of D-APV on spike frequency on the same neurons. APV, 66.99% ± 11.85; wash, 97.67% ± 2.15. N(cells)=7, *, *P*<0.05, paired t-test. **d,** Reduced rheobase current in the presence of M-8324 in the same GABAergic neurons. (**d1**) Sample traces. Scale bars, 100 ms/20 mV. (**d2**) Population data. Pre-drug, 74.29 ± 21.25 pA; M-8324, 62.86 ± 18.22 pA. N(cells)=7. *, *P*<0.05, paired t-test. **e,** M-8324 increased spiking probability measured using a ramp. (**e1**) Sample traces, scale bars, 100 ms/20 mV. (**e2**) Population data. Pre-drug, 53.1% ± 10.79; M-8324, 81.43% ± 14. N(cells)=7, *, *P*<0.05, paired t-test. Data shown as mean±SEM.

Spontaneous spiking is triggered by either synaptic inputs (EPSPs) or fluctuations in membrane potentials ^39, 40^. Under *in vivo* conditions, a large portion of spikes are driven by the preceding EPSPs. This coupling between EPSPs and spikes is termed E-S coupling, and measures the probability of spiking triggered by similar sized EPSPs. E-S coupling is also considered a measure of intrinsic excitability ^22^. We found that the degree of E-S coupling, as measured by the number of spikes generated by five depolarization pulses, was significantly higher in the GABAergic neurons in the presence of M-8324 compared with that with vehicle (Fig. 3a). This enhancement was present in both FS- and RS-neurons (Fig. 3b), suggesting that it occurs in the GABAergic neurons in general. A reversible reduction in the spike numbers triggered by EPSPs was observed with bath perfusion of D-APV (Fig. 3c). In addition, M-8324 significantly shortened the latency to spike compared to the pre-drug baseline (Fig. 3d), consistent with higher excitability. We also found a significant increase in spiking probability by M-8324 using one test pulse (Sup Fig. 3). To exclude the possibility that changes in EPSPs underlies the enhanced E-S coupling, we examined EPSPs in the trials without spikes, in the presence and absence of M-8324. We found no significant difference in EPSP area (Fig. 3e) or waveform (Sup Fig. 4). Altogether, our results indicate a bi-directional modulation of intrinsic excitability by NMDARs in the inhibitory neurons.

**Figure 3.**
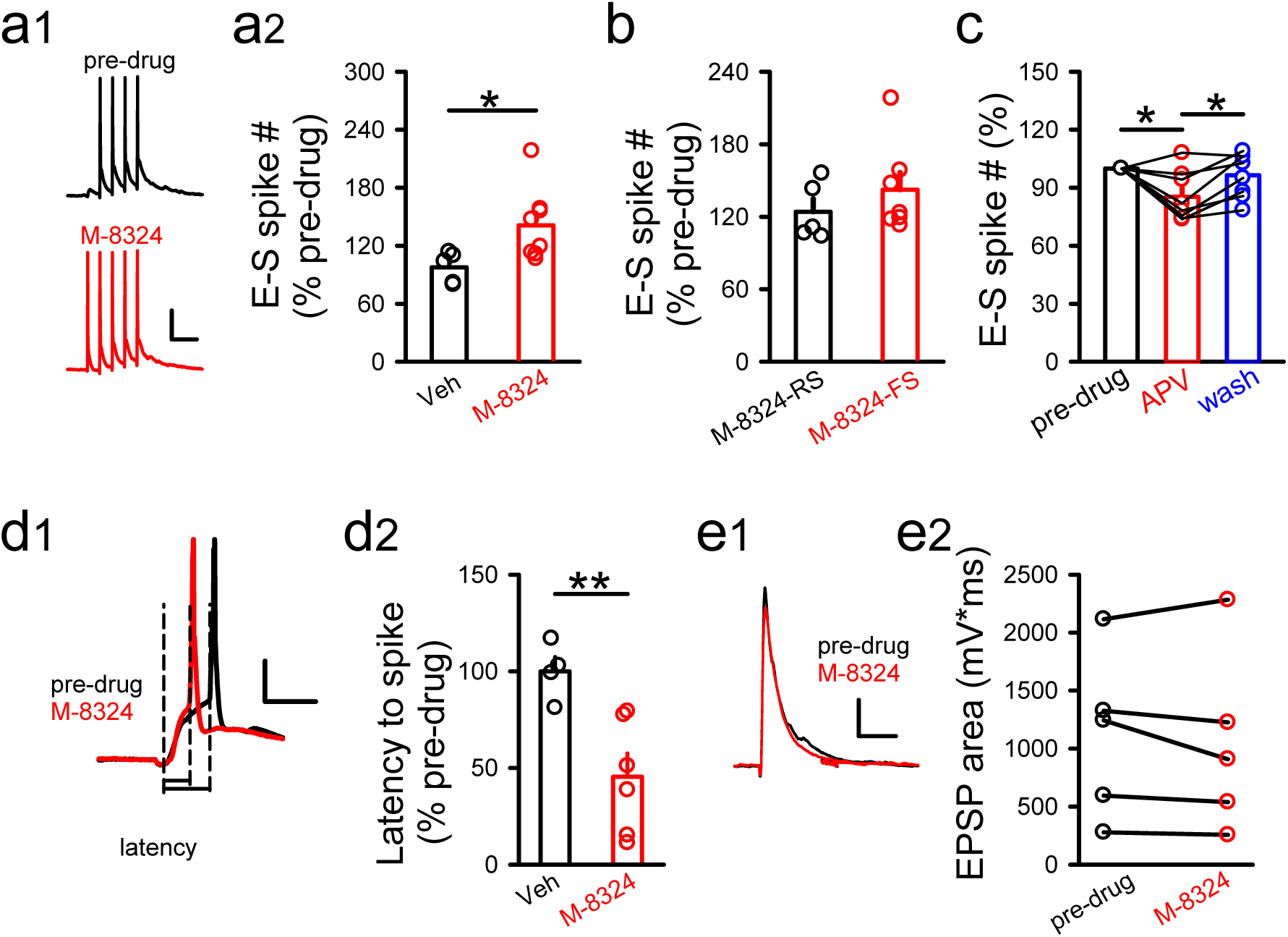
Enhanced E-S coupling by M-8324 in the PFC GABAergic neurons *in vitro*. **a,** Impact of M-8324 on E-S coupling in GABAergic neurons. Sample traces (**a1**) and population data (**a2**). Scale bars, 100 ms/20 mV. Veh, 97.79% ± 7.13, N(cells)=6; M-8324, 141.19% ± 13.27. N(cells)=8, *, *P*<0.05, unpaired t-test. **b,** Impact of M-8324 on E-S coupling in RS- and FS-GABAergic neurons. M-8324(RS), 124.08% ± 10.65, N(cells)=5; M-8324(FS), 142.43% ± 14.13, N(cells)=7. **c,** Impact of D-APV on E-S coupling in the same neurons. APV, 85.36% ± 4.51; wash, 96.45% ± 4.02; both normalized to pre-drug level. N(cells)=8, *, *P*<0.05, paired t-test. **d,** M-8324 shortened the latency to spike. Sample traces (**d1**) and population data (**d2**). Scale bars, 10 ms/20 mV. Veh, 100% ± 7.38, N(cells)=4; M-8324, 45.47% ± 12.03, N(cells)=6. **, *P* <0.01, unpaired t-test. **e,** M-8324 did not affect EPSPs in GABAergic neurons. Sample traces (**e1**) and population data (**e2**). Scale bars, 50 ms/5 mV. EPSP area (mV*ms): 1113 ± 318.9 (pre-drug) and 1043 ± 351.2 (M-8324). N(cells)=5. Data shown as mean±SEM.

### GluN2-subunits have distinct contributions to NMDAR’s modulation of excitability of GABAergic neurons

Distinct contributions of GluN2-NMDAR subunits have been shown for various important NMDAR functions ^2–4^. Thus, we examined the possibility that GluN2-subunits have distinct contributions to the modulation of intrinsic excitability in the GABAergic neurons. To do so, we tested the impacts of modulators selectively targeting different GluN2-containing NMDARs on spontaneous spiking and E-S coupling. Significant increase in spontaneous spiking or E-S coupling induced by M-8324 still occurred in the presence of TCN-201 (10 μM), a GluN2A-selective NMDAR antagonist ^41^ (Fig. 4a). Selective GluN2B-NMDAR antagonist piperidine-18 (1 μM) also did not block increase of spontaneous spiking or E-S coupling induced by M-8324 (Fig. 4b). No enhancement of spontaneous spiking or E-S coupling by M-8324 was observed in the presence of selective GluN2C/2D antagonist NAB-14 (20 μM) (Fig. 4c), while GluN2C/2D-selective NMDAR enhancer CIQ (20 μM) significantly increased spontaneous spiking or E-S coupling (Fig. 4d). The above results suggest that GluN2C/2D-NMDARs play pivotal roles in modulating the intrinsic excitability of GABAergic neurons, while GluN2A- or GluN2B-NMDARs have minimal contributions. To further confirm the contribution of GluN2C/2D-NMDARs, we performed conditional GluN2D knockdown by injecting vGAT-Cre virus into GluN2D conditional knockout GluN2D-Loxp transgenic mice. To confirm the efficacy of this manipulation, we first injected the CMV-Cre virus and found significant reduction in GluN2D protein level in PFC (Fig. 4e). The GABAergic neurons inside the virus-infected regions showed no change in either spontaneous spiking or E-S coupling to application of M-8324 (Fig. 4f), supporting the preferential contribution of GluN2C/2D-NMDARs.

**Figure 4.**
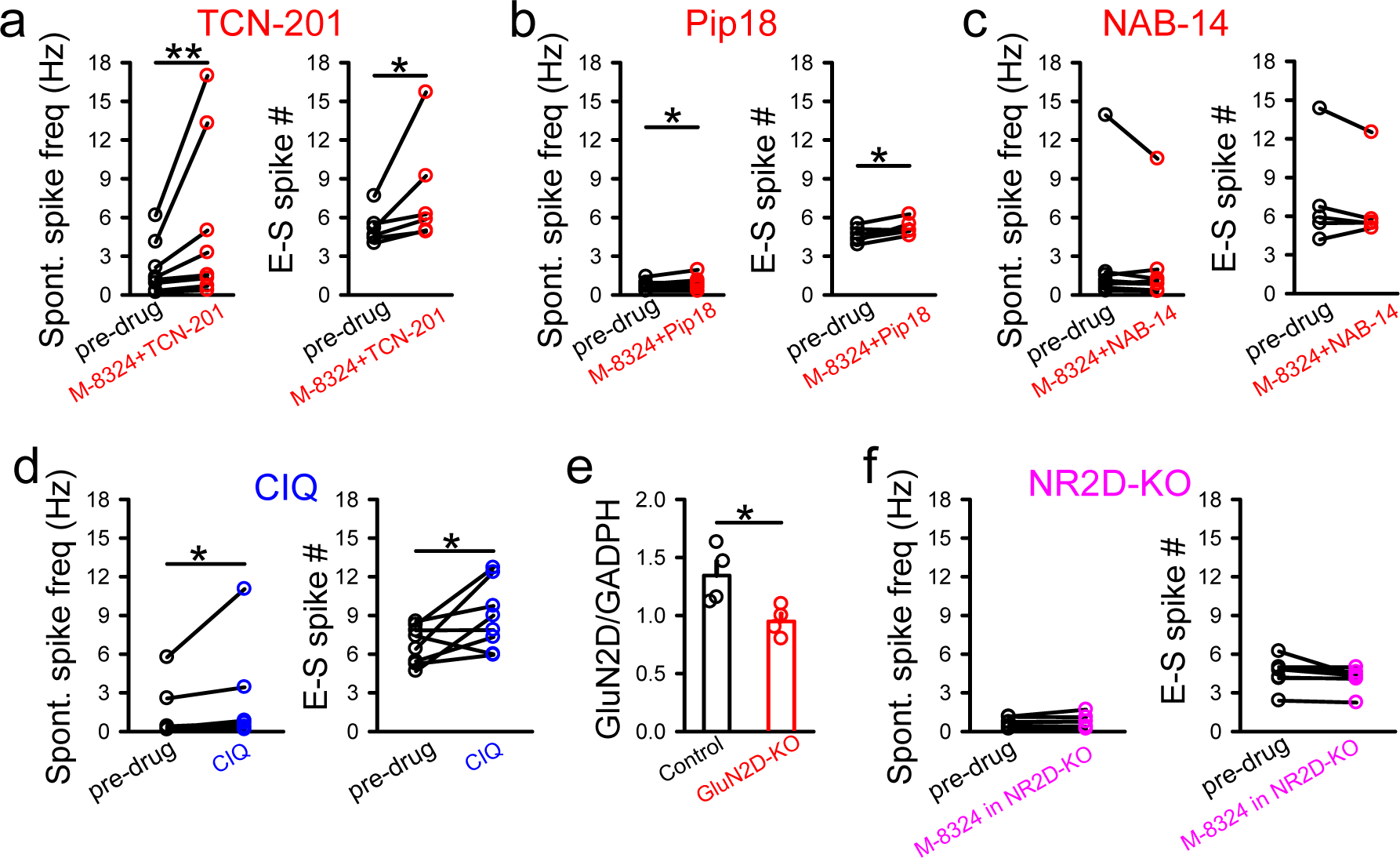
Contributions of GluN2-NMDARs to M-8324’s impact on spontaneous spiking and E-S coupling in GABAergic neurons *in vitro*. **a,** Impact of TCN-201 (10 μM) on M-8324’s effect on spontaneous spiking and E-S coupling. Spontaneous spiking, pre-drug, 2.04±0.73 Hz, M-8324+TCN-201, 5.30±2.23 Hz. N(cells)=8 cells. E-S coupling, pre-drug, 5.22±0.53, M-8324+TCN-201, 7.81±1.70, N(cells)=6. **, *P*< 0.01; *, *P*<0.05, paired t-test. **b,** Impact of Pip18 (1 μM) on M-8324’s effect on spontaneous spiking and E-S coupling. Spontaneous spiking, pre-drug, 0.71±0.12 Hz; M-8324+Pip18, 0.93±0.17 Hz. N(cells)=6. E-S coupling, pre-drug, 4.63±0.24; M-8324+Pip18, 5.21±0.23. N(cells)=6. *, *P*<0.05, paired t-test. **c,** Impact of CIQ (10 μM) on M-8324’s effect on spontaneous spiking and E-S coupling. Spontaneous spiking, pre-drug, 1.22±0.70 Hz; CIQ, 2.26±1.3 Hz. N(cells)=8. E-S coupling, pre- drug, 1.22±0.70; CIQ, 2.26±1.30, N(cells)=8, * *P*<0.05, paired t-test. **d,** Impact of NAB-14 (20 μM) on M-8324’s effect on spontaneous spiking and E-S coupling. Spontaneous spiking, pre-drug, 2.36±1.45 Hz; M-8324+NAB-14, 1.93±1.09 Hz. N(cells)=8. E-S coupling, pre-drug, 7.32±1.80; M-8324+NAB-14, 6.85±1.41. N(cells)=5. *, *P*<0.05, paired t-test. **e,** Potentiation of M-8324 was absent in the inhibitory neurons with GluN2D-NMDAR KO. Spontaneous spiking, pre-drug, 0.69±0.13 Hz; M-8324+NAB-14, 0.75±0.19 Hz. N(cells)=7. E-S coupling, pre-drug, 4.54±0.38, M-8324+NAB-14, 4.22±0.30. N(cells)=8. Data shown as mean±SEM.

### Extrasynaptic GluN2C/D-NMDARs have major contribution to modulating the intrinsic excitability of GABAergic neurons

In the above experiments, the participating NMDARs may be located at either synaptic or extrasynaptic regions of GABAergic neurons. We have shown previously that M-8324 enhances the activity of synaptic NMDARs (Deng, 2020), and here we examined whether it can also enhance the activity of extrasynaptic NMDARs. To do so, we isolated NMDAR responses in the presence of ambient glutamate in the extracellular space ^26, 42, 43^. Bath application of D-APV (100 μM) resulted in a larger reduction in NMDAR responses in the presence of M-8324 than in the vehicle (Fig. 5a), indicating a significant impact of M-8324 on extrasynaptic NMDARs. In addition, NMDAR responses showed larger noise in the presence of M-8324 than in vehicle (data not shown), consistent with larger NMDAR responses ^26^. We also used another widely accepted method to activate extrasynaptic NMDARs, namely puffing NMDA onto neurons ^44^. Both peak and area of NMDAR responses were larger in the presence of M-8324 than in vehicle (Sup Fig. 5), supporting enhanced activation of extrasynaptic NMDARs by M-8324.

To address the question of whether enhancing the activity of extrasynaptic NMDARs may increase the intrinsic excitability of GABAergic neurons, we need to modulate extrasynaptic NMDARs selectively. A NMDAR modulator named EU1794-4 has been suggested to act selectively on the extrasynaptic NMDARs ^45^. We observed significantly larger ambient NMDAR responses in the presence of EU1794-4 than that in the presence of vehicle in the GABAergic neurons (Fig. 5b), suggesting EU1794-4 activates/enhances extrasynaptic NMDARs. For the isolated NMDAR-EPSCs in the GABAergic neurons, there was no significant difference between EU1794-4 and vehicle (Fig. 5c), indicating minimal effect of EU1794-4 on synaptic NMDAR responses. An alternative explanation for the differential impact of EU1794-4 on synaptic vs. extrasynaptic NMDARs is its use-dependence ^45^, which we did not observe using 20 Hz synaptic stimulation (Sup Fig. 6). To further confirm that EU1794-4 selectively binds to the extrasynaptic NMDARs, we synthetized an EU1794-4 compound with a fluorophore attached (see Methods, named EU1794-4-Fluo). This fluorescent EU1794-4 also enhanced extrasynaptic NMDAR responses (Sup. Fig. 7). We then incubated this fluorescent EU1794-4 with primary neuronal culture, and stained the culture with a synapse marker synaptophysin. We found that there was not much overlapping between the fluorescence of these two markers, with the fluorescence of EU1794-4 mostly surrounding that of synaptophysin (Sup. Fig. 8). This result is consistent with EU1794-4 selectively binding to extrasynaptic NMDARs.

We next used EU1794-4 to examine whether extrasynaptic NMDARs play a major role in modulating the intrinsic excitability of GABAergic neurons. We found that EU1794-4 significantly enhanced spontaneous spiking (Fig. 5d), E-S coupling (Fig. 5e, Sup Fig. 9) and current-injection induced spiking (Fig. 5f), indicating a significant contribution of extrasynaptic NMDARs. The increase by EU1794-4 on E-S coupling did not differ between RS- and FS-GABAergic neurons (Sup Fig. 9).

Since GluN2C/2D-NMDARs but not GluN2A- or GluN2B-NMDARs play a prominent role in modulating the intrinsic excitability of the GABAergic neurons (Fig. 4), we examined whether EU1794-4 shows any selectivity on GluN2-NMDARs. Compared to vehicle, NAB-14 abolished changes in holding current caused by bath perfusion of EU1794-4 (Fig. 5g). Considering that it is widely accepted that extrasynaptic NMDARs in the excitatory neurons contain GluN2B subunits ^10, 46^, we further tested whether GluN2B-NMDARs contribute to EU1794-4 activated ambient NMDAR responses. Pip18 did not significantly affect changes in holding current caused by EU1794-4 (Fig. 5h). Furthermore, Pip18 did not affect the increase in spontaneous spiking or E-S coupling in response the presence of NMDA, which enhance the basal activity (Sup Fig. 10). Put together, GluN2C/2D-NMDARs but not GluN2B-NMDARs have significant contributions to EU1794-4’s effect on the extrasynaptic NMDARs. Since the experiments on E-S coupling or spontaneous spiking were done in normal [Mg^2+^] and at -60 mV (Fig. 4), different from the EU1794-4 experiments shown above (at +40 mV; Fig. 5h), we conducted further experiments to confirm that extrasynaptic NMDARs activated under the above physiological conditions contain GluN2C/2D-but not GluN2A- or GluN2B-NMDARs. To do so, slices were bathed in NMDA (3 μM) and changes in holding currents in responses to D-APV were used as a measure of extrasynaptic NMDAR responses, at -60 mV and in 1 mM Mg^2+^. NAB-14 or TCN-201 induced a significantly larger reduction on this holding current comparing to that by Pip18 (1 μM) or DMSO (Fig. 5i), while there was no significant difference between changes induced by Pip18 and DMSO. This result suggests that GluN2C/GluN2D-, GluN2A- but not GluN2B-NMDARs have significant presence at the extrasynaptic locations on the GABAergic neurons.

### Modulation of spiking in GABAergic neurons by NMDAR modulators in vivo

Previous studies, including our own, have shown that NMDAR inhibitors lead to reduced spiking of inhibitory neurons *in vivo* ^34, 47^. We recently showed that NMDAR-PAMs elevate spiking of the inhibitory neurons in the auditory cortex *in vivo* (Deng, 2020). Our current results indicate that the excitability of GABAergic neurons is modulated bi-directionally by NMDARs, especially those at the extrasynaptic locations. To test whether this modulation also occurs *in vivo* and involves extrasynaptic NMDARs, we first examined the impact of M-8324 on neuronal spiking in the PFC *in vivo* since PFC is a key brain region for many critical brain functions (such as working memory and executive functions) and various psychiatric disorders (including schizophrenia and depression). By using the well-established criteria to distinguish between excitatory and inhibitory neurons (Sup Fig. 11), we observed significantly enhanced spiking in the inhibitory neurons and reduced spiking in the excitatory neurons after infusion of M-8324 (Sup Fig. 11), consistent with our previous finding in the auditory cortex ^33^. The reduced spiking level in excitatory neurons is likely caused by the enhanced spiking in the inhibitory neurons ^26, 33, 48^. To further investigate whether modulation the extrasynaptic NMDARs may contribute to neuronal spiking, *in vivo* cell-attached recording was used to record the sound-induced spiking and spontaneous spiking in PV-neurons in the primary auditory cortex (A1; Fig. 6a). These neurons in PV-Cre mice were confirmed positive for parvalbumin based on their responses to opto-stimulation injected with ChR2 virus (see Methods) (Fig. 6a-c). In addition, their spike waveforms were characteristic for PV-neurons (Fig. 6c). Both sound-evoked and spontaneous spiking frequency (Fig. 6d-f) were significantly and reversibly enhanced after local infusion of EU1794-4 (Fig. 6a). In comparison, significant and reversible reduction in both sound-evoked and spontaneous spiking was observed in the pyramidal/excitatory neurons in A1 (Fig. 6g-i). Taken together, we have observed enhanced spiking in the inhibitory neurons to application of EU1794-4 *in vivo*, in a manner similar to that *in vitro*, suggesting that activation of extrasynaptic NMDAR *in vivo* also significantly enhances the activity of inhibitory neurons.

**Figure 5.**
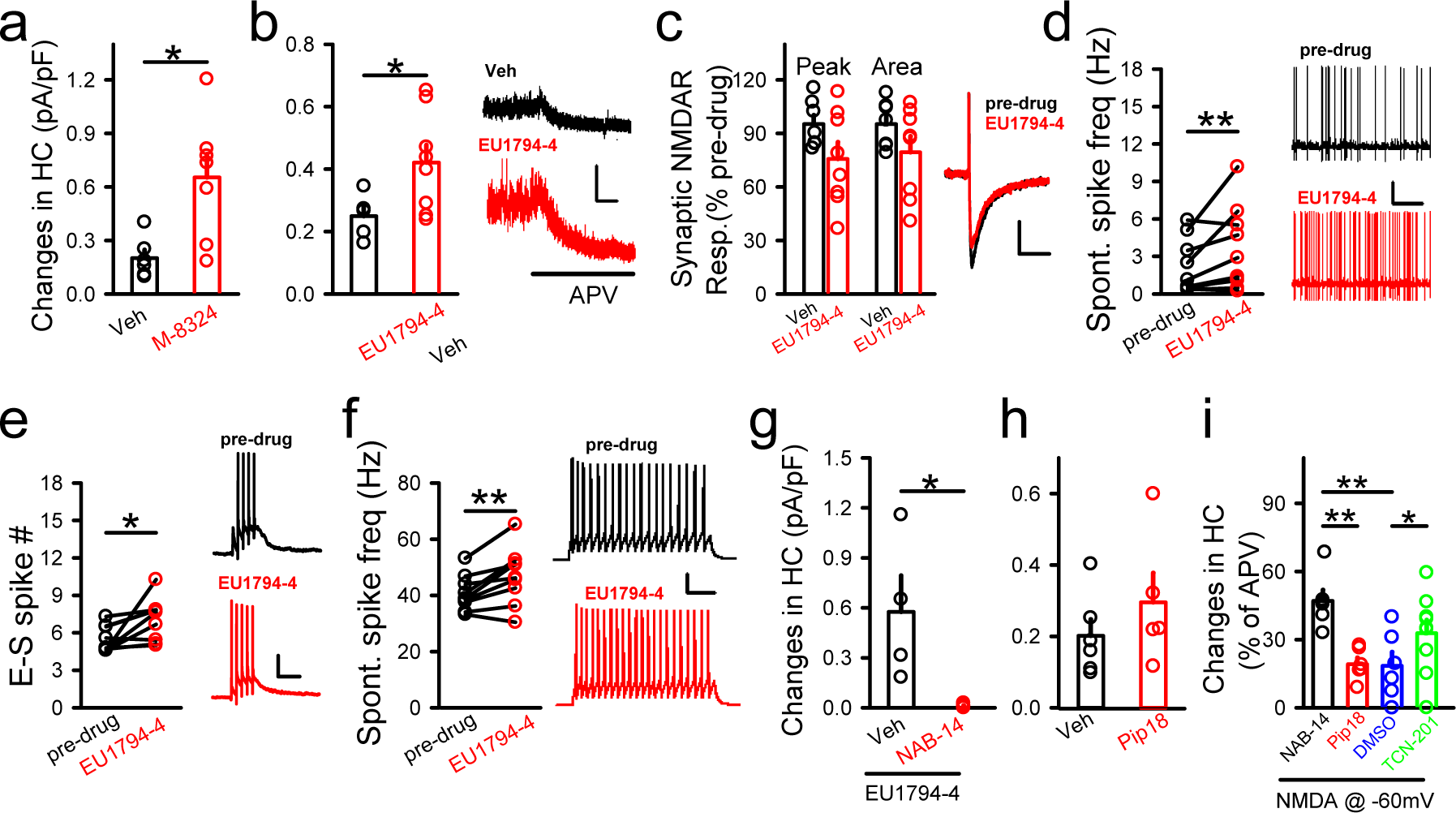
Contribution of extrasynaptic NMDARs to the intrinsic excitability of GABAergic neurons. **a,** Ambient NMDAR responses in the presence of M-8324 or vehicle. Veh, 0.20 ± 0.046 pA/pF, N(cells)=6; M-8324, 0.65 ± 0.13 pA/pF. N(cells)=7. *, *P*<0.05, unpaired t-test. **b,** Ambient NMDAR responses in the presence of EU1794-4 (30 μM) or vehicle, with sample traces on the right. Veh, 0.24 ± 0.03 pA/pF, N(cells)=5; EU1794-4, 0.42 ± 0.05 pA/pF, N(cells) =8. Scale bars, 10 s/50 pA. *, *P*<0.05, unpaired t-test. **c,** EU1794-4 on evoked NMDAR-EPSCs, compared to vehicle, with sample traces on the right. Scale bars, 100 ms/20 pA. Peak amplitude, Veh, 95.22% ± 5.01; EU1794-4, 75.71% ± 9.43; area, Veh, 95.24% ± 4.95; EU1794-4, 79.46% ± 8.92. N(cells)=7 (Veh), 8 (EU1794-4). **d,** EU1794-4 on spontaneous spike frequency normalized to baseline, with sample traces on the right. Scale bars, 10 s/20 mV. Pre-drug, 1.84±0.56 Hz; EU1794-4, 3.07± 0.88 Hz. N(cells)=9. **, *P*<0.01, paired t-test. **e,** EU1794-4 on E-S coupling, with sample traces on the right. Scale bars, 200 ms/20 mV. Pre-drug, 5.47±0.35; EU1794-4, 7.05±0.56. N(cells)=11. *, *P*<0.05, paired t-test. **f,** EU1794-4 on spikes triggered by current injections, with sample traces on the right. Scale bars, 500 ms/10 mV. Pre-drug, 40.76 ±2.12 Hz; EU1794-4, 46.76 ±3.35 Hz. N(cells)=9. **, *P*<0.01, unpaired t-test. **g,** Impact of NAB-14 (20 μM) on EU1794-4 (300 μM)-induced changes in holding current at +40 mV. Veh, 0.57 ± 0.21 pA/pF, N(cells)=4; NAB-14, 0.01 ± 0.0001 pA/pF, N(cells)=5. *, *P*< 0.05, unpaired t-test. **h,** Impact of Pip18 on the ambient NMDAR responses in GABAergic neurons. Veh, 0.20 ± 0.046 pA/pF, N(cells)=6; Pip18, 0.29 ± 0.082 pA/pF, N(cells)=5. **i,** Impacts of various GluN2-NMDAR antagonists on isolated NMDAR responses in the presence of NMDA (3 μM) at -60 mV. Changes in holding current to bath perfusion of antagonists were normalized to changes in holding currents after subsequent D-APV application. NAB-14, 46.94% ± 5.89, N=5; Pip18, 19.20% ± 2.82, N(cells)=6; DMSO, 15.81% ± 5.78, N(cells)=7; TCN-201, 32.80% ± 5.83, *, *P*<0.05, **, *P*<0.01, unpaired t-test. Data shown as mean±SEM.

**Figure 6.**
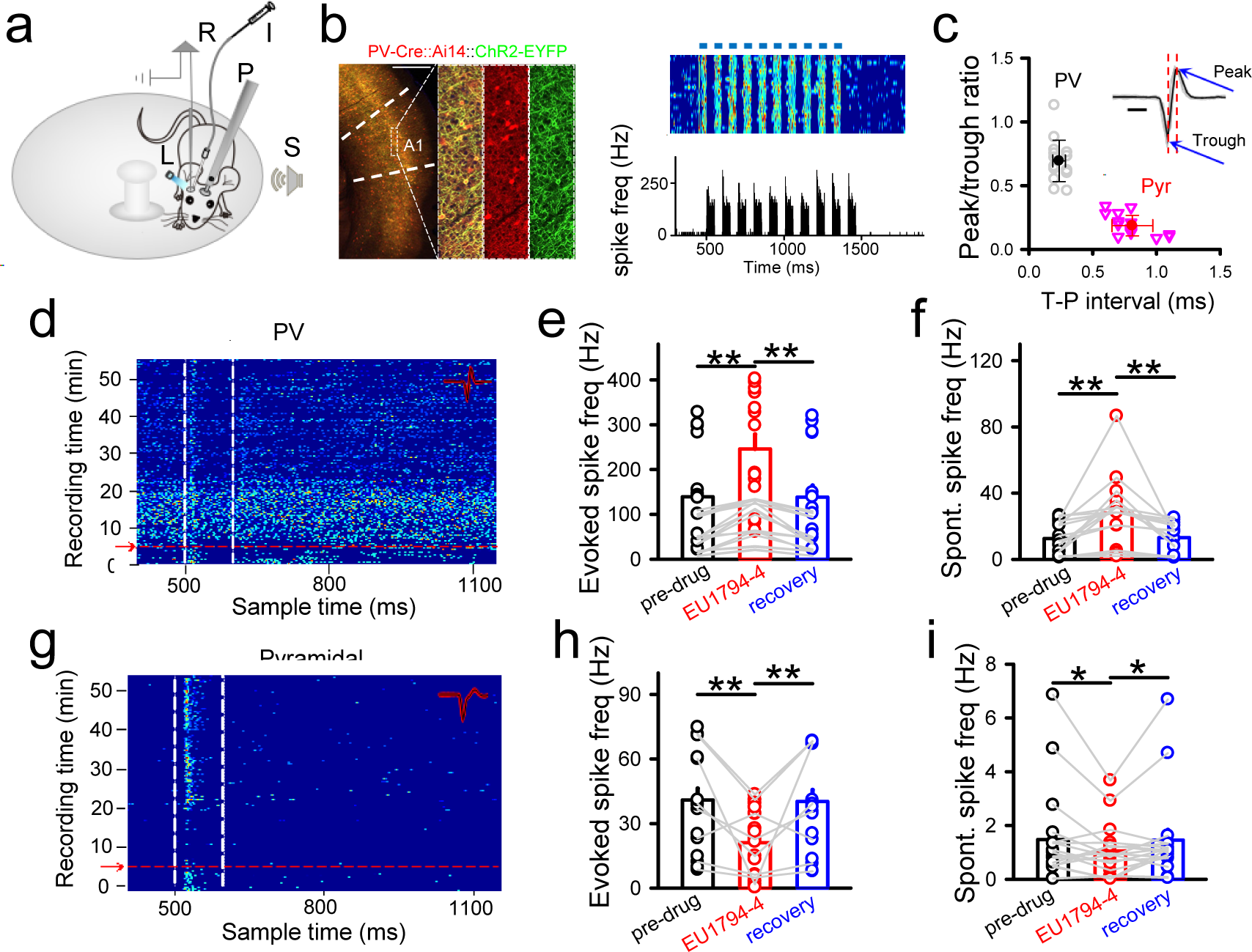
Impact of EU1794-4 on neuronal spiking *in vivo*. **a,** Experimental setup. A mouse was head-fixed via a headpost (P) but could run freely on a rotatable plate. Sound (S) was applied to one ear and patch recording (R) was performed in the contralateral A1. Blue Light (L) and drug infusion tube (I) were put next to recording site of A1. **b,** (Left) Confocal images show tdTomato-labeled PV-neurons (red) and expression of ChR2-YFP (green) in a representative slice. Scale bar, 500 μm. (Right top) Raster plot of spikes in an example PV-neuron to pulses of blue LED light stimulation (blue bars, 50 ms each pulse). (Right bottom) Corresponding post-stimulus spike time histogram. **c,** Peak/trough amplitude ratio plotted against trough-to-peak interval of spike waveform. Each data point for an individual neuron. Solid symbols represent mean ± SD. Inset: spike waveforms of an example PV-neuron. Black traces showing 20 superimposed spikes. Red dotted vertical lines mark the timing of trough and peak. Blue arrows point to peak and trough. Peak/trough ratio: Pyr, 0.19 ± 0.08; PV, 0.69 ± 0.16. Trough-peak interval: Pyr, 0.81 ± 0.16; PV, 0.24 ± 0.05; Scale bar, 0.5 ms. **d-f,** (**d**) Noise-evoked responses (raster plots) of PV-neurons in A1 before and after drug injection (red arrow). Dashed lines indicate the onset and offset of acoustic stimulation. Inset: 20 randomly selected superimposed spike waveforms. Noise-evoked (**e**) and spontaneous (**f**) spike frequency of recorded PV-neurons before, 15 min and recovery after EU1794-4 infusion. N=16 cells. **, *P* < 0.01, paired t-test. **g-i,** Similar to (**d-f**) but for Pyramidal neurons. N(cells)=18. **, *P* < 0.01, *, *P* < 0.05, paired t-test. All data shown as mean±SEM.

## Discussion

In this study, we have revealed important contributions of extrasynaptic NMDARs to the intrinsic excitability of inhibitory/GABAergic neurons: (1) NMDARs bidirectionally modulates this excitability in that activation enhances while inhibition reduces it; (2) GluN2C/2D-containing NMDAR subunits have a significant contribution to this modulation; (3) extrasynaptic NMDARs play a critical role in this modulation. These new findings have far-reaching implications for the contributions of NMDARs to both physiological functions and pathology of brain diseases.

Mostly based on studies on the excitatory neurons, extrasynaptic NMDARs have been shown to play a few distinct functions: (1) their excessive activation leads to synapse loss and neuronal loss especially in diseases such as stroke ^1, 4^; (2) their activation can modulate neuronal excitability in developing excitatory neurons ^17, 49^; (3) their suppression by ketamine (especially via GluN2B-contaning NMDARs) mediates enhanced glutamatergic synaptic transmission and is proposed to underlie the anti-depressant effect of ketamine ^50–52^. Although inhibition of NMDARs on the inhibitory neurons has also been suggested to mediate ketamine’s anti-depression effect, the location of these NMDARs has not been examined ^53–55^. Here we provide strong evidence that extrasynaptic NMDARs play an important role in modulating the intrinsic excitability of inhibitory neurons in a bi-directional manner. This finding emphasizes their role in the second-to-second routine functions of inhibitory neurons in contrast to their much-emphasized role in synaptic plasticity. There are some evidences that NMDARs are required for the induction of synaptic plasticity in the inhibitory neurons ^6^, but the subcellular localization of these NMDARs is unclear.

### Subunit composition and subcellular localization of NMDARs participate in the modulation of intrinsic excitability

We have previously characterized M-8324 and showed that M-8324 has higher potency than GNE-8324 and only potentiates synaptic NMDAR responses on the inhibitory neurons (Deng, Masri et al. 2020). We do not have information on the subunit selectivity of M-8324, although it is likely similar to that of GNE-8324. Since extrasynaptic NMDARs are mainly activated by the extracellular or ambient glutamate under the physiological conditions, their activation may be subjected to modulations distinct from that for synaptic NMDARs, such as less activity-dependence and no input-specificity ^56^. This point can be appreciated for depolarization-induced or spontaneous spiking in that this modulation likely involves most or all extrasynaptic NMDARs on an entire neuron and is affected by extracellular glutamate concentration surrounding the entire neuron. For E-S coupling, however, the situation can be complex since synaptically released glutamate may affect the extrasynaptic NMDARs surrounding the active synapses in the inhibitory neurons ^26, 57^.

NMDARs with different GluN2 subunits have been shown to be present at distinct synaptic localizations and serve distinct functions, although this conclusion has been debated ^2, 10, 46^. More specially, extrasynaptic NMDARs are shown to be enriched with GluN2B-, GluN2C- or GluN2D-NMDARs in the excitatory neurons, while GluN2C/2D-containing NMDARs are present at extrasynaptic regions in the inhibitory neurons mostly found in young animals ^27, 42, 58^. Since their activation requires spillover of synaptic glutamate or elevated extracellular glutamate, extrasynaptic NMDARs on the excitatory neurons show more prominent activation under pathological condition or at hyperactivity brain state ^46^. Pharmacological evidence suggest that GluN2C/2D-contaning NMDARs can modulate the activity of both GABAergic neurons and excitatory neurons ^29, 58, 59^. Our results strongly support this conclusion on the inhibitory neurons and further indicate the absence of GluN2B-, GluN2A-containing NMDARs at the extrasynaptic regions in the inhibitory neurons, distinct from the excitatory neurons. This lack of extrasynaptic GluN2A-, GluN2B-containing NMDARs is also consistent with our finding that modulation of intrinsic excitability occurs in the presence of normal extracellular Mg^2+^ concentration and near resting membrane potentials, and consistent with the low sensitivity of GluN2C/2D-NMDARs to Mg^2+^ block at negative membrane potentials ^2, 60^. Since FS and Non-FS interneurons do not show significant difference on NMDAR’s impacts on the intrinsic excitability, it is thus likely that extrasynaptic NMDARs on these neurons do not differ substantially, such as on GluN2-subunit composition. This is in contrast to the less contribution of synaptic NMDARs in FS than non-FS inhibitory neurons ^43, 61^, and further supports potentially different contributions between synaptic and extrasynaptic NMDARs to the inhibitory neuronal functions.

With extracellular glutamate level playing a major role in their activation, this mode of modulation of extrasynaptic NMDARs is distinct from the activity/input-dependent modulations typically occurring during synaptic plasticity. The modulation of intrinsic excitability by extrasynaptic NMDARs is influenced by the level of extracellular/ambient glutamate, and/or the density and/or composition of extrasynaptic NMDARs. Significant alterations in the extracellular/ambient glutamate levels occur during development and under pathological conditions including epilepsy, stroke and Alzheimer’s disease ^58, 62–64^. The density/activity of extrasynaptic NMDARs has been shown to be altered in Alzheimer’s disease, Huntington diseases and stroke ^65, 66^. In theory, changes in the subunit composition of extrasynaptic NMDARs will affect the activation of these receptors, such as switching from GluN2C/2D-NMADRs to GluN2A/2B-NMDARs. How the above factors may interact to influence the activation of extrasynaptic NMDARs to affect the intrinsic excitability of inhibitory neurons and brain functions requires further exploration.

### Pathological contribution of extrasynaptic NMDARs in schizophrenia

Our finding of modulation of intrinsic excitability by extrasynaptic NMDAR may also have important implications for the hypofunction of NMDARs in schizophrenia. Evidence from both human postmortem and animal models indicate that reduced activity/expression of NMDARs in the inhibitory neurons (especially PV-neurons) play key roles in the pathogenesis and/or pathology of schizophrenia ^6, 11, 12, 67^. Due to the low presence of synaptic NMDARs on certain subtypes of adult inhibitory neurons (especially PV-neurons) ^61, 68–70^, it is not apparent how this hypofunction might occur or causing pathology (Moreau and Kullmann 2013). Our current findings provide suggest a potential solution to puzzle: reduced presence/activity of extrasynaptic NMDARs mediates this hypofunction in inhibitory neurons (especially PV-neurons), leading to reduced neuronal excitability and imbalanced excitation/inhibition in the brain which is generally assumed to be critical for the pathogenesis/pathology in schizophrenia. An implication of this model is that alterations and consequences of reduced NMDAR function occur in an input-non-specific manner and hence may affect a wider range of functions than those associated with selective inputs onto inhibitory neurons ^71^.

### Potential mechanisms underlying the modulation of intrinsic excitability by NMDARs

We here showed similar impacts of NMDAR modulators on three measurements of intrinsic excitability (spontaneous spiking, depolarization-induced spiking and E-S coupling) in the inhibitory neurons. Previous studies in the excitatory neurons have demonstrated an interaction between Kv4.2 and NMDARs. Kv4.2 channels are present at extrasynaptic sites in the excitatory neurons ^72, 73^, and they may interact with both synaptic and extrasynaptic NMDARs. Activation of extrasynaptic NMDARs results in the dephosphorylation of Kv4.2 and increased neuronal excitability, and this process is mediated by protein kinase A, protein kinase C, extracellular signal-regulated kinase and CaMKII ^74–77^. Recent evidence showed that the GSK3β directly interacted with Kv4.2 via Ser-616 in medium spiny neurons ^78^. Thus, a potential modulation of IA/Kv4.2 channels to mediate the impacts of NMDARs on intrinsic excitability in the inhibitory neurons is worthy of further examination. Other mechanisms mediating NMDAR modulation of excitability might also exist, such as via NMDAR activity-dependent regulation of HCN and Akt/mTOR signaling ^79, 80^. In addition, inhibition/ablation of extrasynaptic NMDARs increases the neuronal excitability via repression of Kv2.1 or Kv1.1 ^81, 82^, suggesting that linking NMDAR activity with that of K+ channels might be a general mechanism in modulating excitability. Whether this mechanism may differ between excitatory and inhibitory neurons is worthy of further exploration.

### NMDAR-PAMs

We have used NMDAR-PAMs to enhance the activation of NMDARs in order to understand their contributions to the excitability of inhibitory neurons. Comparing to NMDAR antagonists, there is no upper limit for the enhancement induced by PAMs and hence their impact can be revealed as long as some residual activation of NMDARs exists. This strategy is especially useful for *in vitro* analysis since it is likely that the level of NMDAR activation is low in brain slices compared to *in vivo*. Although a large number of NMDAR-PAMs have been developed ^37^, for treating diseases with reduced inhibitory neuron activity NMDAR-PAMs that selectively enhance the activity of inhibitory neurons (such as GNE-8324 and M-8324) (Hackos, 2016; Yao, 2018; Deng, 2020) will likely be more useful by avoiding potential counteraction and excitotoxicity due to enhanced activation of excitatory neurons (Hanson, et al., 2020).

In conclusion, our findings provide strong support for NMDARs having important impacts on the second-by-second neuronal functions in the inhibitory neurons by modulating the intrinsic excitability bidirectionally. Since excitability influence many critical functions of neurons and brain states, we suggest that NMDARs may play wider and more important roles in the inhibitory neurons than we have previously recognized.

## Acknowledgments

We thank Chang Ji for performing the RT-PCR experiment. This work was supported by Shenzhen Bay Laboratory (SZBL2019062801003), Youth Program of the National Natural Science Foundation of China (No.82004469), Shenzhen-Hong Kong Institute of Brain Science-Shenzhen Fundamental Research Institutions (2019SHIBS0004), Opening Operation Program of Key Laboratory of Acupuncture and Moxibustion of Traditional Chinese Medicine in Guangdong (NO.2017B030314143),

## Author Contributions

L.L.Y, experiment design, data collection and analysis, discussion and writing; R. Y, experiment design, data collection and analysis, discussion; X.Y.M, experiment design, data collection and analysis; H.F.L, experiment design, data collection and analysis; D.D, experiment design, data collection and analysis, discussion and writing; Y.J.C, discussion and writing; S.Y, discussion and writing; T.P, discussion and writing; T.Y, discussion and writing; F.X.L, discussion and writing; N.G.X, discussion and writing; Q.Z, experiment design, data collection and analysis, discussion and writing.

## Data and code availability

All data generated and codes created during the current study are available from the lead contact upon reasonable request.

## Declaration of Interests

The authors declare no competing interests.

---

**Sup Figure 1.**
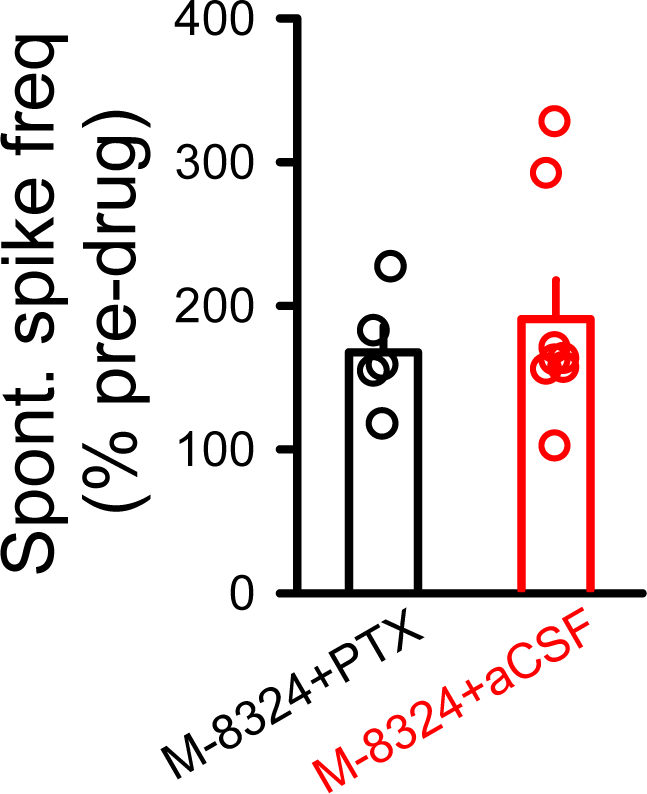
Picrotoxin (100 μM) did not affect M-8324’s effect on spontaneous spiking in GABAergic neurons. M-8324 (PTX), 167.47% ± 18.01, N(cells)=5; M-8324 (aCSF), 190.75% ± 27.13, N(cells)=8.

**Sup Figure 2.**
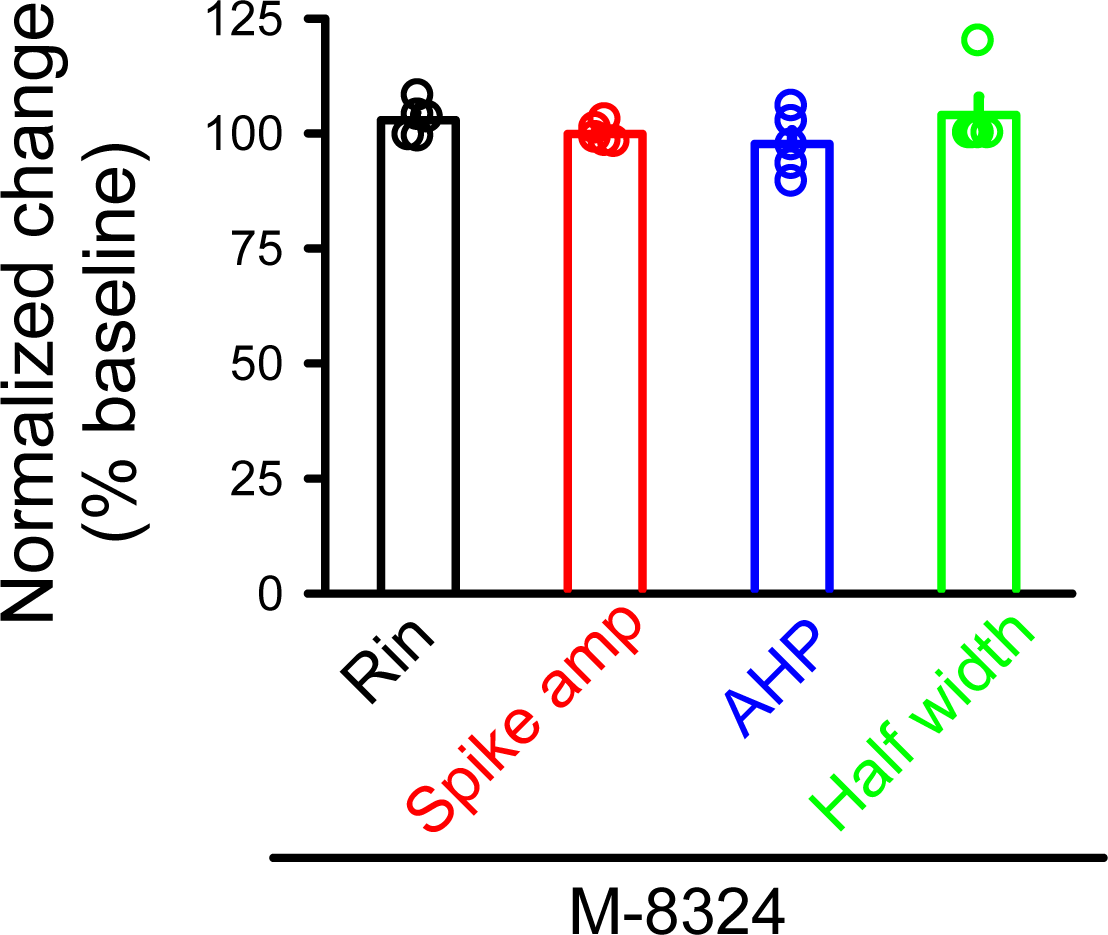
M-8324 did not affect basic neuronal properties in GABAergic neurons. **A,** Input resistance (101.60% ± 2.21), N(cells)=5; **B**, spike amplitude (99.90% ± 0.91). **C,** afterhyperpolarization (97.72% ± 0.91). **D,** spike half width (105.41% ±3.56).

**Sup Figure 3.**
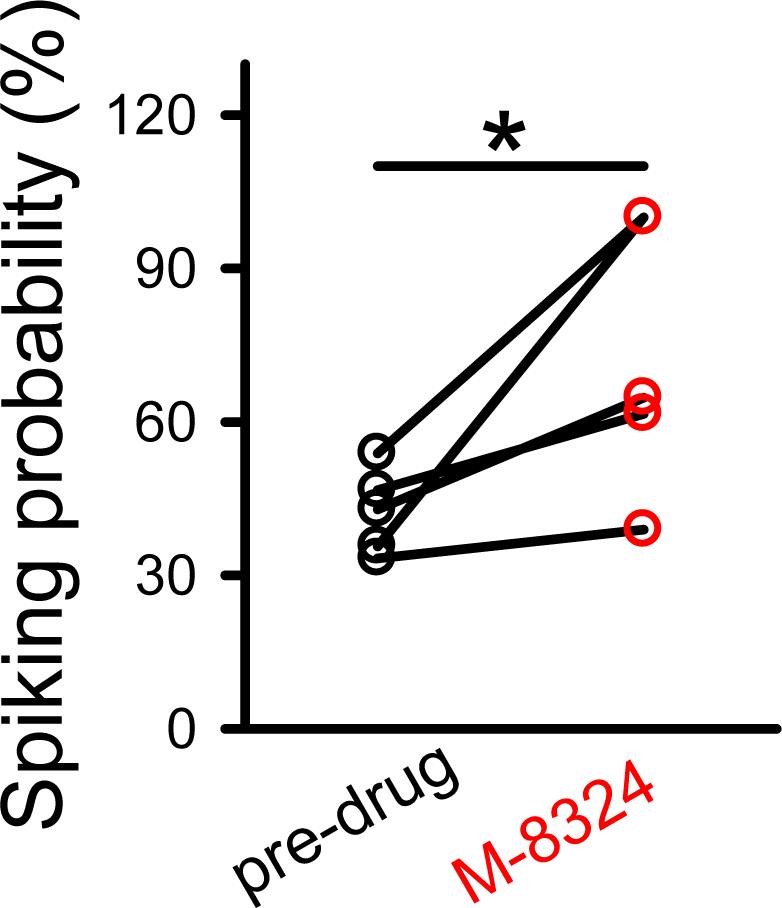
M-8324’s impact on the probability of single EPSP-induced spike. Pre-drug, 42.48% ± 3.71; M-8324, 73.02% ± 11.87. N(cells)=5, *, *P*<0.05, paired t-test.

**Sup Figure 4.**
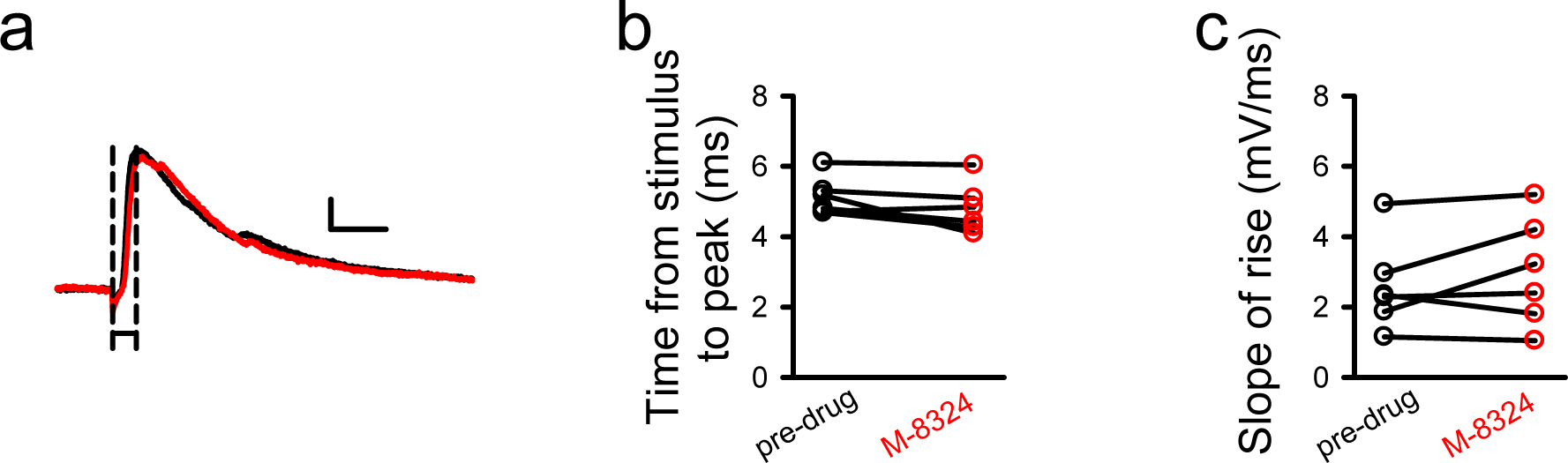
No significant impact of M-8324 on EPSPs. **(a)** Sample traces of EPSP showing the parameter of measurement. Scale bars, 10 ms/2 mV. Neither time to peak (**b**) nor slope of rise time (**c**) was altered by M-8324. N(cells)=5.

**Sup Figure 5.**
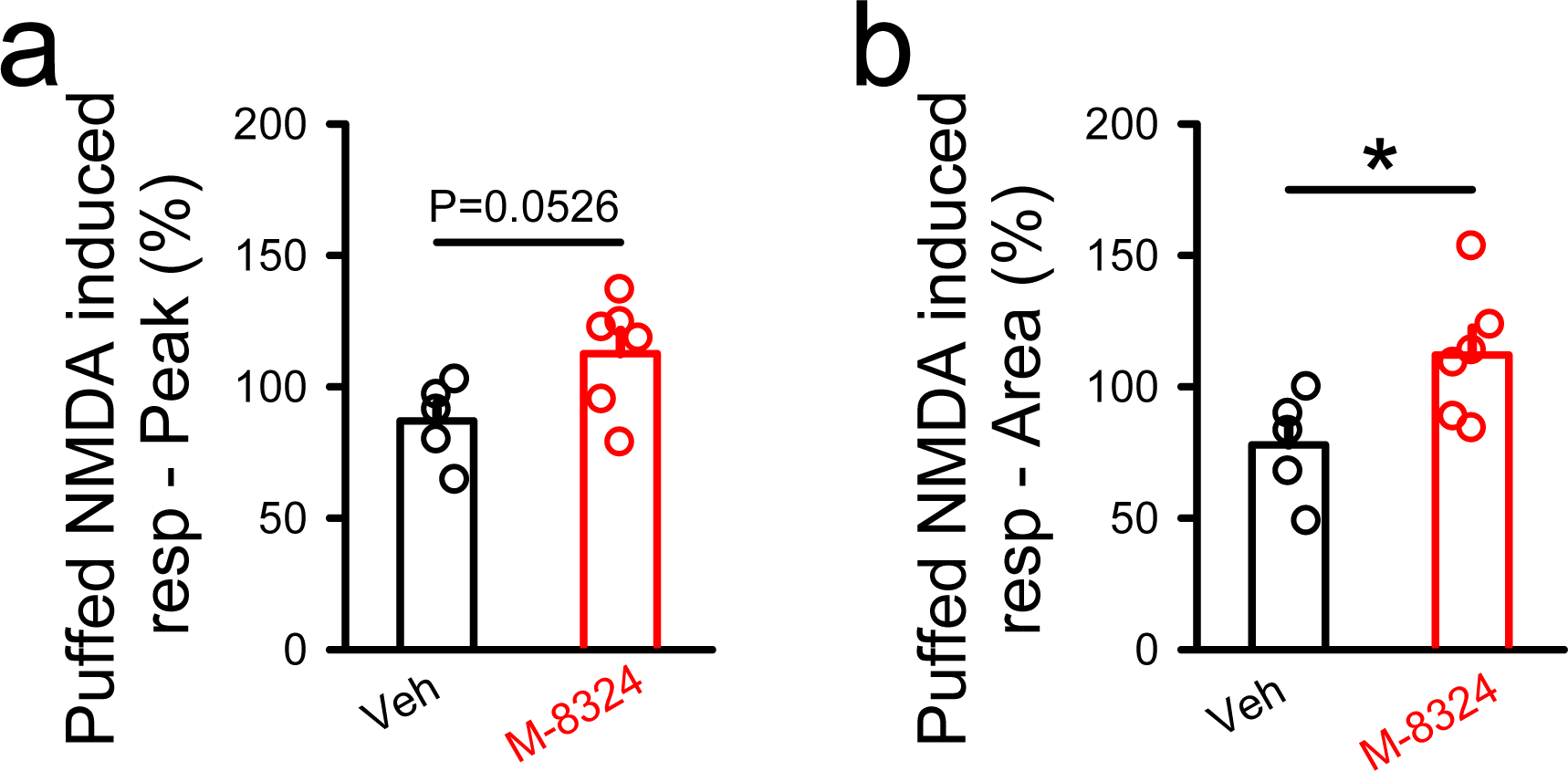
M-8324 increases extrasynaptic NMDAR responses induced by puffed NMDA. Peak (**a**) and area (**b**) of puffed NMDA-induced response in the presence of M-8324 or Veh. Veh, Peak, 87%±6.77; Area,112.62%±8.79, N(cells)=5; M-8324, Peak, 77.81%±8.92; Area, 112.03%±10.28, N(cells)=6. *, *P*<0.05, unpaired t-test.

**Sup Figure 6.**
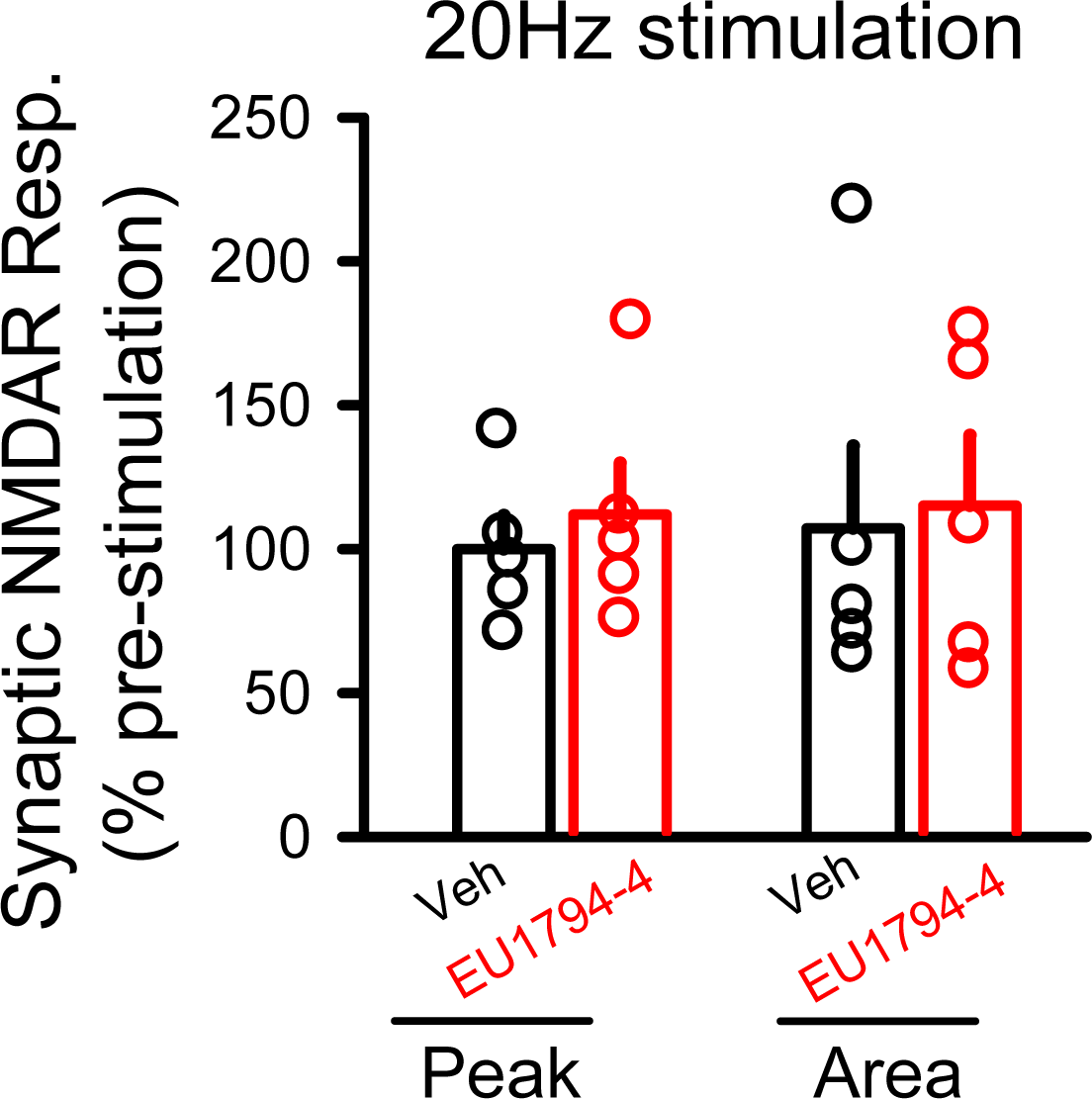
Activity-dependence of EU1794-4 on NMDAR-EPSCs. Neither peak nor area was altered after 20 Hz stimulation in the presence of EU1794-4 (30 μM). Veh, Peak, 99.98%±11.8; Area, 107.18%±28.77, N(cells)=5; EU1794-4, Peak, 112.07%±17.87’ Area, 115.12%±24.42, N(cells)=5.

**Sup Figure 7.**
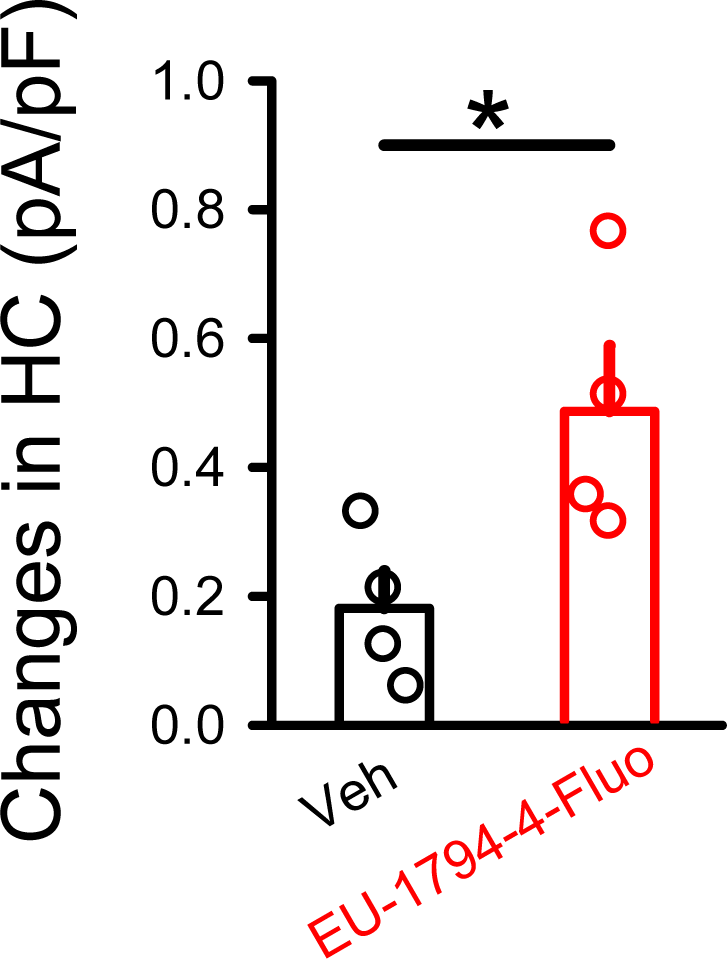
Fluorophore-attached EU1794-4 (EU1794-4-Fluo) increases extrasynaptic NMDAR responses. Ambient NMDAR responses were significantly larger in the presence of EU1794-4-Fluo (300 μM), comparing to the Veh. Veh, 0.18 ± 0.11, N(cells)=4; EU1794-4-Fluo, 0.48 ± 0.10, N(cells)=4. ##, *P*<0.01, unpaired t-test.

**Sup Figure 8.**
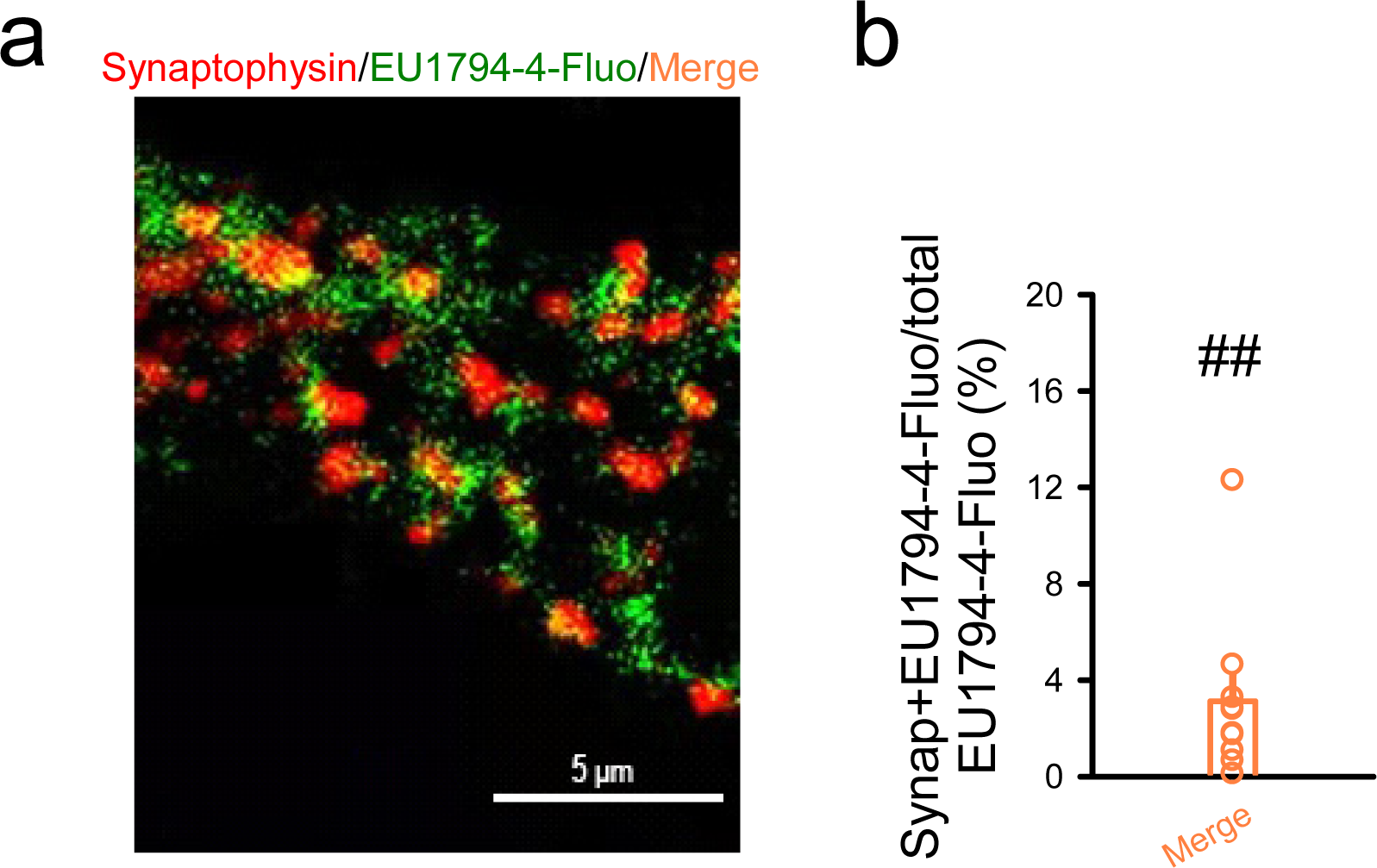
Staining of EU1794-4-Fluo with synaptophysin on cultured neurons. (**a**) Representative images of synaptophysin (Red), EU1794-4-Fluo (Green) and merge. (**b**) Percentage of synaptophysin + EU1794-4-Fluo/total EU1794-4-Fluo. 3.11% ± 1.10, N=10. ##, *P* <0.01, paired t-test.

**Sup Figure 9.**
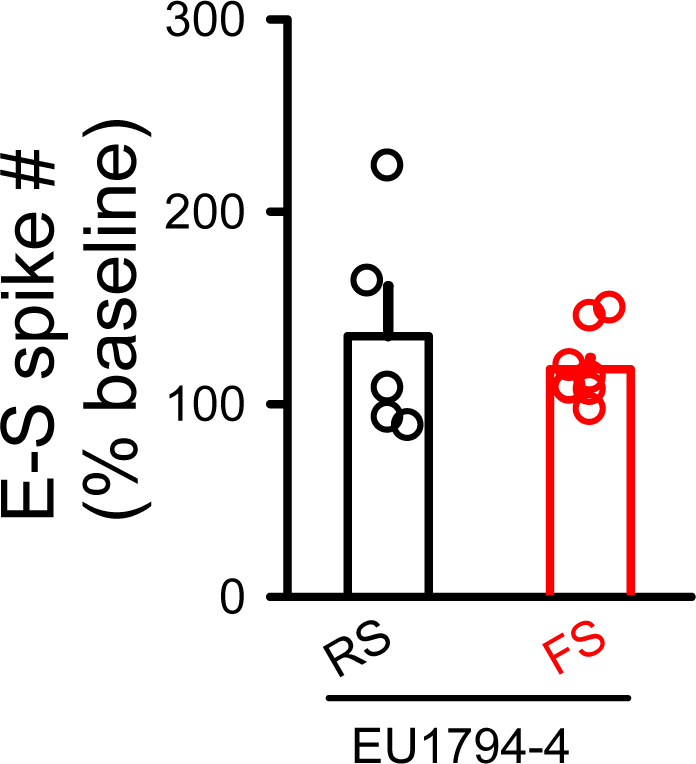
The impact of EU1794-4 on E-S coupling did not differ between RS- and FS- inhibitory neurons. RS,135.38% ± 25.82, N(cells)=5; FS, 118.14% ± 5.94, N(cells)=9.

**Sup Figure 10.**
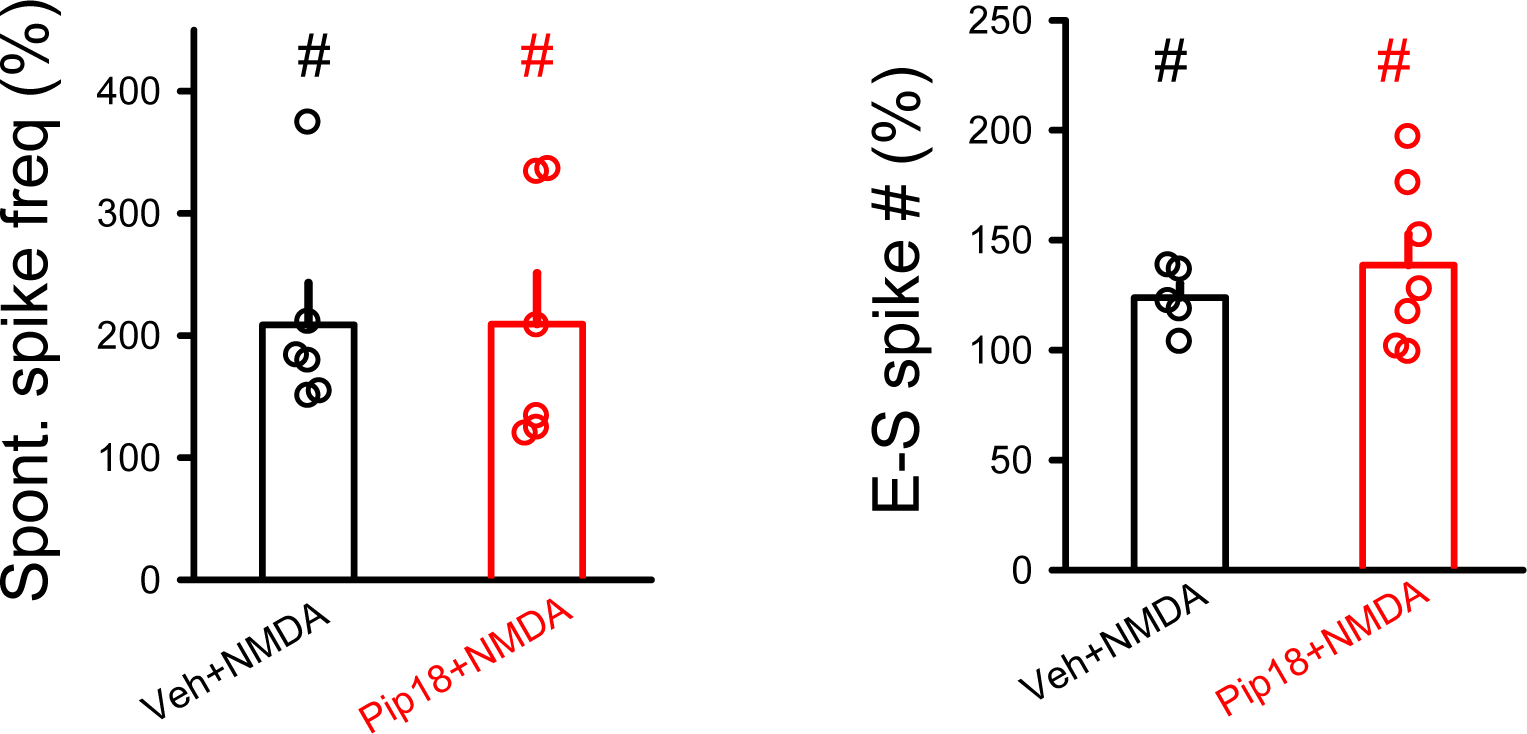
Pip18 did not affect the increase of spontaneous spiking or E-S coupling induced by NMDA. Spontaneous spiking experiment, Veh+NMDA, 208.67%±34.26, N(cells)=6; Pip18+NMDA, 209.21%±41.82, N(cells)=6, #, *P*<0.05, paired t-test, comparing to that in the pre-drug. E-S coupling experiment, Veh+NMDA, 123.88%±6.37, N(cells)=5; Pip18+NMDA, 138.60%±14.20, N(cells)=8, #, *P*<0.05, paired t-test, comparing to the pre-drug.

**Sup Figure 11.**
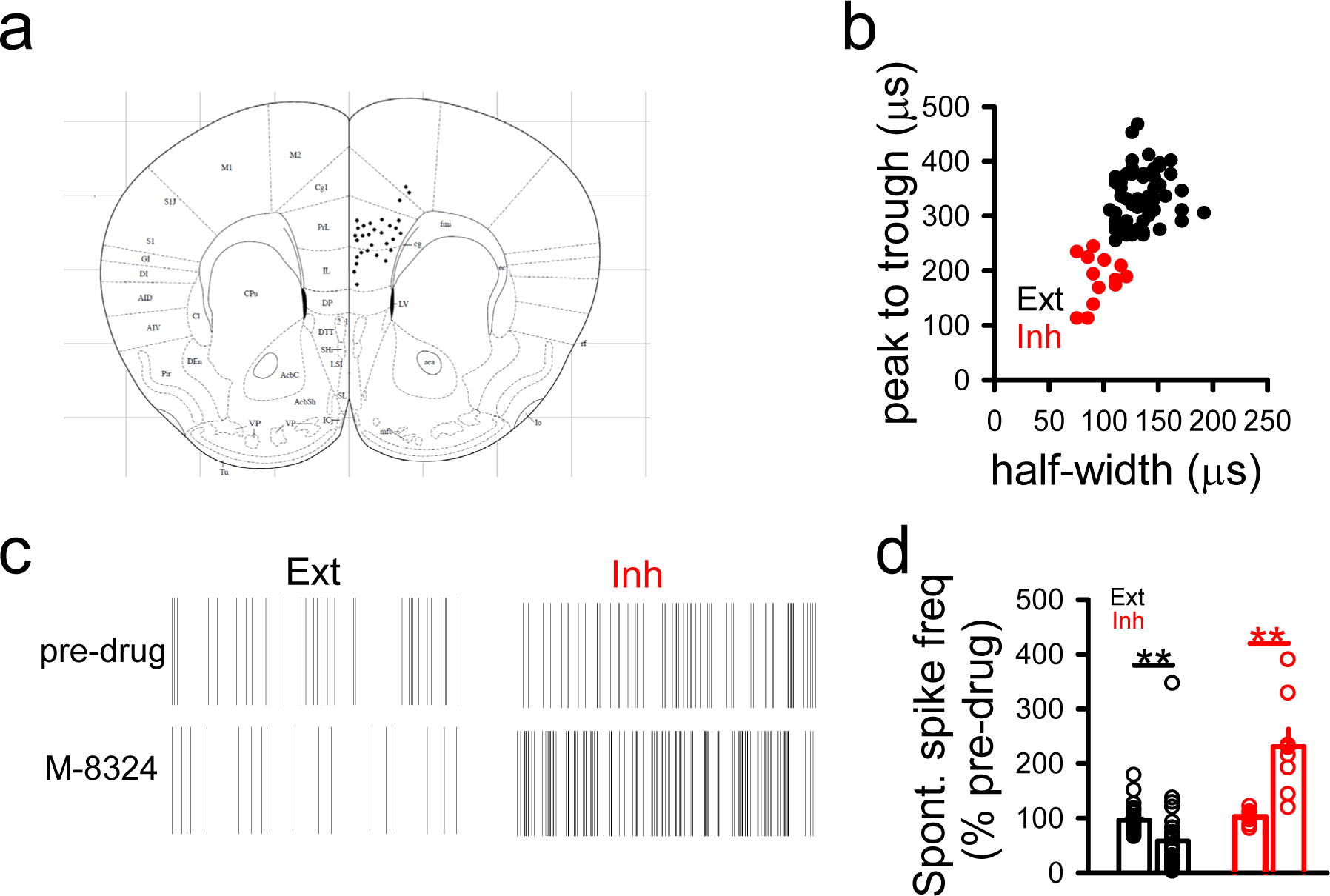
Modulation of spontaneous spiking in PFC neurons in *vivo*. **(a)** The location of recording electrodes was in the PFC. **(b)** Criteria for distinguishing fast spiking neurons (inhibitory neurons, Inh) and non-fast spiking neurons (excitatory neurons, Ext) based on their spike waveforms. (**c)** Sample raster plots showing effect of M-8324 (ICV, 100 μM) on spontaneous spiking of inhibitory neurons. Total duration, 10 sec. **(d)** Quantification of NMDAR modulator effects on spontaneous spiking in excitatory and inhibitory neurons. For excitatory neurons, 58.01% ±10.39, N =35 units/5 mice (M-8324, ICV, 100 μM). For inhibitory neurons, 230.76% ±31.80. N=9 units/5 mice.

## Notes

### Competing Interest Statement

The authors have declared no competing interest.

